# Prediction of allosteric sites and mediating interactions through bond-to-bond propensities

**DOI:** 10.1101/056275

**Authors:** B.R.C. Amor, M.T. Schaub, S.N. Yaliraki, M. Barahona

## Abstract

Allosteric regulation is central to many biochemical processes. Allosteric sites provide a target to fine-tune protein activity, yet we lack computational methods to predict them. Here, we present an efficient graph-theoretical approach for identifying allosteric sites and the mediating interactions that connect them to the active site. Using an atomistic graph with edges weighted by covalent and non-covalent bond energies, we obtain a bond-to-bond propensity that quantifies the effect of instantaneous bond fluctuations propagating through the protein. We use this propensity to detect the sites and communication pathways most strongly linked to the active site, assessing their significance through quantile regression and comparison against a reference set of 100 generic proteins. We exemplify our method in detail with three well-studied allosteric proteins: caspase-1, CheY, and h-Ras, correctly predicting the location of the allosteric site and identifying key allosteric interactions. Consistent prediction of allosteric sites is then attained in a further set of 17 proteins known to exhibit allostery. Because our propensity measure runs in almost linear time, it offers a scalable approach to high-throughput searches for candidate allosteric sites.

## I. INTRODUCTION

Allostery is a key molecular mechanism underpinning control and modulation in a variety of cellular processes [1, 2]. Allosteric effects are those induced on the main functional site of a biomolecule by the binding of an effector at a distant site, e.g., the binding of a cofactor modulating the catalytic rate of an enzyme [3]. Despite the importance of such processes, there is still a lack of understanding as to how the interactions at the allosteric site propagate across the protein and affect the active site. In this paper, we present a graph-theoretic approach that uses atomistic structural data to identify allosteric sites in proteins, as well as bonds and residues involved in this propagation. By defining an edge-to-edge transfer function, which can be understood as a Green’s function in the edge space of the protein graph, we compute a bond-to-bond propensity that captures the effect induced on any bond of the molecule by the propagation of perturbations stemming from bonds at the active site. This propensity can be computed efficiently to predict allosteric sites and key bonds which are prominently involved in mediating the allosteric propagation.

The growing realisation that all proteins exhibit innate dynamic behaviour [4, 5] and the discovery of allosteric effects in single domain proteins [6] have reaffirmed the ubiquitousness of this form of regulation; potentially, any protein could be allosteric [7]. This fact opens up important experimental directions: drugs targeted at allosteric sites could offer improved specificity and control compared to traditional drugs that bind at the active site [3]. Efficient methods able to identify putative allosteric sites are therefore of great current interest [8]. To date, computational approaches to finding allosteric sites have involved statistical coupling analysis [9], molecular dynamics [10–12], machine learning [13], and normal mode analysis [14]. For a comprehensive review see Ref. [15].

Classic thermodynamic models of allostery (such as the Monod-Wyman-Changeux (MWC) [16] and Koshland-Nemethy-Filmer (KNF) models [17]) were formulated to explain cooperativity in multimeric proteins in terms of conformational transitions in a protein landscape [11, 18]. Although such models reproduce broad experimental features (e.g. the sigmoidal binding curves), they offer little insight into the molecular mechanisms driving and defining the underlying conformational transitions. Attempts to identify the specific residues involved in allosteric transitions have led to the idea of allosteric pathways, which aim to describe the routes through which an excitation propagates through the protein [9, 19, 20]. Indeed, recent experimental [21, 22] and computational [23–26] work has shown that energy flow in globular proteins is anisotropic. Some of these studies have connected this anisotropy to the allosteric properties of the protein [22, 26]. Our work builds on this line of research and aims at finding allosteric sites by using graph-theoretical techniques to quantify efficiently the propagation of perturbations through a protein structure described in atomistic detail. In [22] the authors find that internal energy flow in albumin is anisotropic, and that this flow is altered by binding of an allosteric ligand. Here, we also find that the propagation of perturbations internally is anisotropic. However, we use the term ‘allosteric’ in a more specific way, to describe locations distant from the active site where a perturbation can have a functional effect on the active site. The identification of such distant sites and the pathways connecting them to the active-site, has become an area of considerable interest [12, 27, 28].

The connection between the behaviour of a diffusion process (e.g., a random walk) on a network and the vibrational dynamics of that network is well established in the biophysical literature [29, 30]. Previous network-based methods for protein structure analysis have made use of shortest path calculations [31], community detection algorithms [32], and random walks on networks [33]. However, such methods almost universally used *coarsegrained* protein descriptions at the level of residues, i.e., they are based on residue-residue interaction networks (RRINs) [34] that neglect atomistic detail. Although methods that use molecular dynamics simulations to derive edge weights for RRINs from the crosscorrelations of residue fluctuations have yielded interesting results [35, 36], such approaches are computationally costly. Furthermore, Ribeiro and Ortiz have recently shown that RRINs are critically dependent on the chosen cut-off distance, and that using *energy-weighted* networks that include the covalent interactions of the backbone is crucial for correctly identifying signal propagation pathways [37, 38]. Our findings below show that efficient methodologies which can exploit the physico-chemical detail of atomistic, energy-weighted protein networks can lead to enhanced identification of allosteric sites and relevant individual mediating interactions in a number of important cases.

Our analysis starts by building an atomistic graph model of the protein: nodes are atoms and (weighted) edges represent individual bonds, with weights given by energies from interatomic potentials. The graph includes both covalent bonds and weak, non-covalent bonds (hydrogen bonds, salt bridges, hydrophobic tethers and electrostatic interactions). Details of the construction of the graph are given in Section IVE and in Refs. [39, 40]. The resulting all-atom graph is analysed using the edge-to-edge transfer matrix *M*, which is akin to a discrete Green’s function in the *edge space* of the graph and has been recently introduced in Ref. [41] to study non-local edge coupling in graphs. In this paper, we derive a new, alternative interpretation of the matrix *M* and show that it provides a means to extracting the level of influence that the fluctuations of an edge have on any other edge of the graph (for detailed mathematical derivations, see Materials and Methods, Section IV A 1 and *SI*). We use this notion to calculate the *propensity* of each bond, Π_*b*_, i.e., a measure of how strongly bond *b* is coupled to the active site through the atomistic graph. Because allosteric effects are reflected on induced changes in weak bonds, yet mediated through the whole protein network, our bond-to-bond formalism provides a natural way of uncovering how the long-range correlations between bonds contribute to allosteric signalling. Crucially, recent algorithmic developments [42, 43] allow these computations to be carried out in *almost linear time* (in the number of edges). Therefore, in contrast to most other computational approaches, our method is easily scalable to large systems with tens of thousands of atoms.

To establish if a bond has a high propensity Π_*b*_, and to detect important bonds (and residues), we use quantile regression to compare each bond to the ensemble of bonds within the protein at a similar geometric distance from the active-site (described in Materials and Methods, Section IV B). Quantile regression (QR) [44] is a robust statistical technique previously employed in medicine [45], ecology [46] and econometrics [47]. We additionally confirm our findings by computing the statistical significance of the bond propensity against a reference set of 100 representative proteins randomly drawn from the Structural Classification of Proteins (SCOP) database (see Section II D). This reference set provides us with a pre-computed structural bootstrap against which any protein can be tested to detect statistically significant bonds, further reducing the computational cost of our method.

In Sections II A—II C, we showcase our procedure through the detailed analysis of three important allosteric proteins: caspase-1, CheY, and h-Ras. In each case, given structural data and the location of the known active site, we correctly predict the location of the allosteric site and uncover communication pathways between both sites. Each of the three examples serves to highlight particular aspects of the method. In the case of caspase-1, comparison of our results with those obtained using coarse-grained residue-residue interactions networks (RRINs) shows that incorporating atomistic physico-chemical detail can indeed be necessary for the reliable identification of the allosteric site. In the case of CheY, we illustrate how further information can be gained by incorporating dynamic data from ensembles of NMR structures: the variance of the propensity across the NMR ensemble reveals residues involved in allosteric signalling which cannot be identified from the static X-ray crystal structure alone. In the case of h-Ras, our method shows that signal propagation between the active and allosteric sites is crucially dependent on the interaction between the protein and specific structural water molecules. Having demonstrated the insight into allosteric mechanisms offered by our method, we then evaluate it against a test set with a further 17 allosteric proteins (see Section IIE). We find that the bond-to-bond propensity is a good predictor of a site’s allosteric propensity, suggesting it could be used to guide efforts in structure-based discovery of drugs as allosteric effectors.

## II. RESULTS

### A. Identification of the allosteric site and functional residues in caspase-1

Our first example is caspase-1, an allosteric protein of great importance in apoptotic processes [40]. Caspase-1 is a tetramer composed of two asymmetric dimers, each containing one active site. Using the PDB atomic structure (PDB: 2HBQ), we constructed an atomistic, energy-weighted graph representation of the protein based on interaction potentials, as described in Section IV E [39, 40].

In order to quantify how strongly each bond is coupled to the active site, we calculate the propensities Π_*b*_ for all bonds in the protein, as given by Eq. (8). We also aggregate the bond propensities for each residue to obtain the residue score Π_*R*_, as given by Eq. (9). To rank bonds and residues according to their significance, we compute the corresponding quantile scores *p_b_* and *p_R_*, respectively, obtained via quantile regression as in Eq. (14). These quantile scores allow us to establish which bonds (and residues) have high propensity values as compared to bonds (or residues) at the same distance from the active site in the protein (Fig. 1a and 1c).

**FIG. 1.**
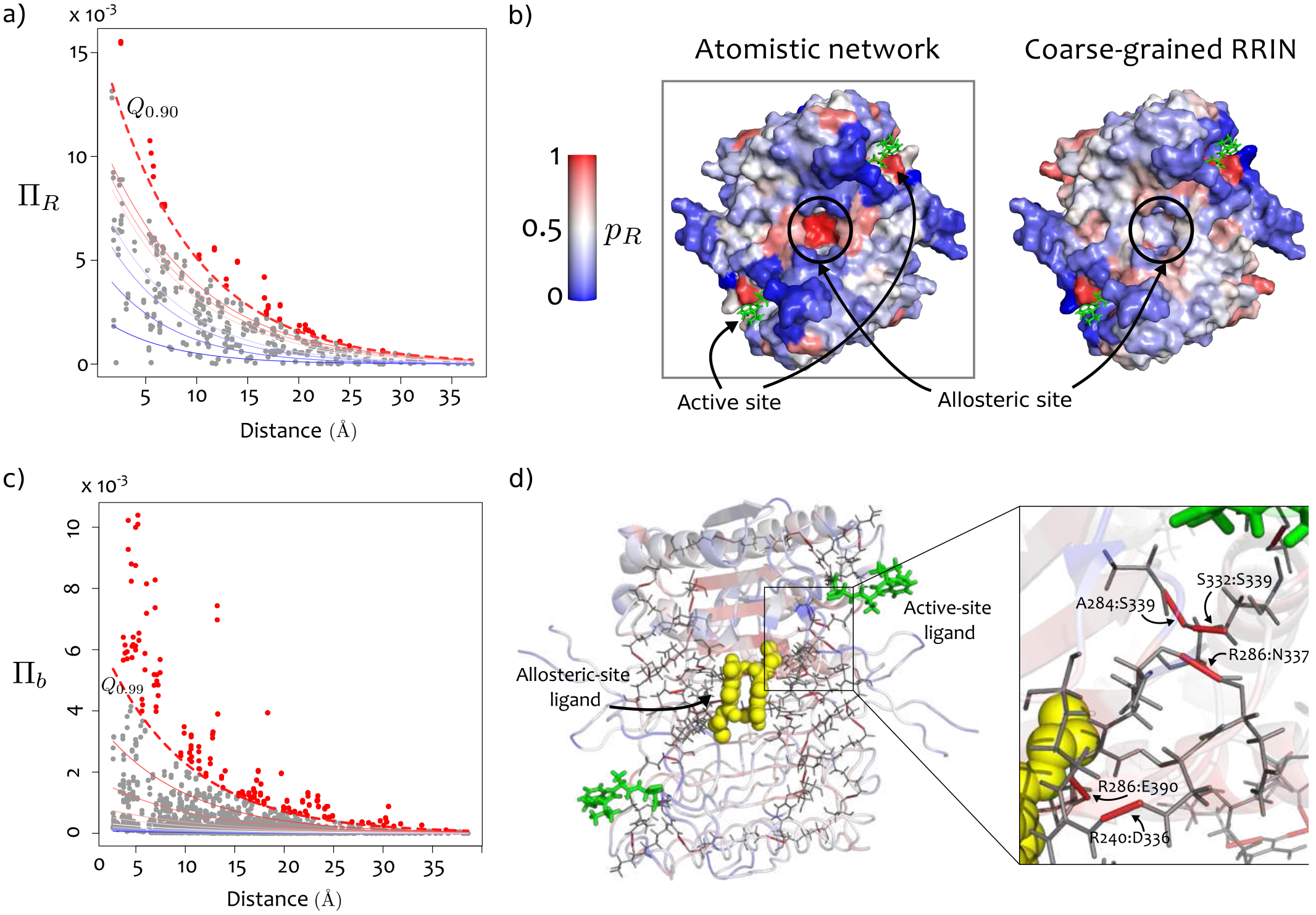
Bond-to-bond propensities identify the allosteric site and atomistic pathway in caspase-1. (a) The propensities of all residues Π_*R*_ are plotted against their distance from the active site. The lines correspond to the quantile regression estimates for the *p*-th quantiles *Q_p_*, with *p* = 0.1, 0.2,…, 0.8, 0.9. The dashed red line indicates the *Q*_0.90_ cut-off used for identifying important residues. (b) The quantile scores *p_R_* for each residue are mapped onto the surface of caspase-1. The active-site ligand is shown in green. The allosteric binding site is identified as a hot-spot of high propensity. When a coarsegrained residue-residue interaction network with cut-off of 6Å is used (right), the allosteric binding site is not identified. (c) The propensities of bonds Π_*b*_ are plotted against their distance from the active site with the *Q*_0.99_ quantile indicated by the dashed line. (d) High quantile score bonds (*p_b_* ≥ 0.99) are shown on the structure. Bonds between R286:E390, R240:D336, R286:N337, A284:S332, and S332:S339 have large quantile scores and form contiguous pathways between the active and allosteric sites. The active site ligand is shown in green and the allosteric ligand is shown as yellow spheres.

Our method finds a hot spot of residues with high quantile scores in a cavity at the dimer-dimer interface (Fig. 1b left). This site has been previously identified by Scheer *et al.* as the binding site for a small molecule inhibitor of caspase-1 [48]. Table I shows that the allosteric residues, i.e., residues within 3.5Å of the allosteric inhibitor, have significantly higher propensities than non-allosteric residues (Wilcoxon rank sum, *p* < 0.0005). Residues E390, S332 and R286, which have been found to belong to a hydrogen bond network between the active and allosteric sites [48], have respectively the 3rd, 13th, and 15th highest quantile scores of the 260 residues in each dimer of caspase-1.

Making use of the physico-chemical detail afforded by our atomistic description, we find the bonds with high propensity that lie on communication pathways connecting the allosteric site to the active-site ligand. Concentrating on the top quantile *p_b_* ≥ 0.99 (Fig. 1c), the two interactions in the salt bridges between residues E390 and R286 have quantile scores of 0.996 and 0.990, and their combined propensity gives this salt bridge the highest quantile score in the protein. It is known that these salt-bridges are directly disrupted by the allosteric inhibitor [48]. In addition, our method reveals other important bonds lying between the active and allosteric sites (Fig. 1d), including hydrogen bonds between Arg240:Asp336 (*p_b_* = 0.999), S332:S339 (*p_b_* = 0.996), R286:N337 (*p_b_* = 0.992), and A284:S332 (*p_b_* = 0.990). Bonds in this pathway have previously been identified by Datta *et al* as being functionally important: the corresponding alanine mutations cause 230-fold (R286A), 130fold (E390A), 3.7-fold (S332A) and 6.7-fold (S339A) reductions in catalytic efficiency [48].

The atomistic detail is important for the outcome of the analysis. If instead of employing an all-atom graph description, we carry out the same calculations on a coarse-grained residue-residue interaction network (RRIN) [31, 33] with cut-off radius of 6 Å, the allosteric site of caspase-1 is no longer identified as a hot spot (Fig. 1b right) and the allosteric residues do not have significantly higher propensity compared to other residues (Wilcoxon rank sum, *p* = 0.5399). The results obtained with RRINs are in general dependent on the cut-off radius used. For caspase-1, the allosteric site is not detected in RRINs with cut-off radii of 6 Å, 7 Å and 8 Å. The allosteric site is found to be significant at 10Å, but the signal is still considerably weaker than when using the atomistic network (Table S6). These findings highlight that while an atomistic model of the protein structure may not always be needed, it can indeed be important for the detection of allosteric effects in proteins; in this case, the strength of the pair of salt bridges formed by E390 and E286, which is crucial for the allosteric communication in caspase-1, is not captured by RRINs. Other recent results have similarly demonstrated the importance of both covalent bonds and hydrogen bonds to signal transmission within proteins [38]. Yet in other cases (e.g., CheY in the following section), this level of physico-chemical detail seems to be less important, and RRINs are able to capture allosteric communication. An extended, indepth analysis of the results obtained with all-atom networks and RRINs for a variety of proteins and cut-off radii can be found in the SI (Section 6).

### B. Uncovering allosteric communication pathways in CheY

#### 1. Identification of the phosphorylation site of CheY

CheY is a key protein in bacterial chemotaxis. When CheY binds to the flagellar motor switch protein (FliM), it causes a change in the rotation direction of the flagellar motor, thus regulating the tumbling rate of *E. coli*. This regulation is achieved through a post-translational modification of CheY: phosphorylation of CheY at the distant residue D57 increases its affinity for FliM, making this an interesting example of a single-domain allosteric protein.

**TABLE I.** Quantile scores for the propensities of residues within 3.5Å of the allosteric site of caspase-1 computed from the atomistic graph and from a residue-based network (RRIN) with cut-off radius of 6 Å. The average quantile scores of allosteric residues 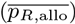 and non-allosteric residues 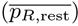 are also presented.

| Residue | $p_R$ (Atomistic network) | | $p_R$ (RRIN) | |
| --- | --- | --- | --- | --- |
|  | Dimer 1 | Dimer 2 | Dimer 1 | Dimer 2 |
| R240 | 0.772 | 0.734 | 0.562 | 0.562 |
| L258 | 0.394 | 0.408 | 0.168 | 0.168 |
| N259 | 0.828 | 0.832 | 0.324 | 0.324 |
| F262 | 0.654 | 0.652 | 0.464 | 0.464 |
| R286 | 0.938 | 0.928 | 0.838 | 0.838 |
| C331 | 0.634 | 0.646 | 0.724 | 0.724 |
| P335 | 0.206 | 0.196 | 0.450 | 0.450 |
| E390 | 0.990 | 0.992 | 0.318 | 0.318 |
| R391 | 0.982 | 0.984 | 0.258 | 0.258 |
| $\overline{p_R^{allo}}$ | 0.711 | 0.708 | 0.4567 | 0.4567 |
| $\overline{p_R^{rest}}$ | 0.481 | 0.492 | 0.4793 | 0.4789 |

Following the same procedure, we calculated the propensity of each bond and residue (relative to the FliM binding site) in fully activated CheY (PDB ID: 1F4V) bound to Mg^2+^, BeF_3_ and FliM. We identify a number of hot-spot surface residues with high quantile scores (Fig. 2a), including the phosphorylation site, D57 (*p_R_* = 0.96). Again, residues in the allosteric site (< 3.5 Å from the phosphorylation site) have higher average quantile score than non-allosteric residues 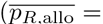 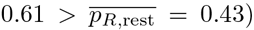), and four of the seven residues in the allosteric site have high quantile scores, *p_R_* ≥ 0.9 (Table II). In addition, we find a number of previously unidentified distant surfaces with high quantile scores (Fig. 2a), which could correspond to putative (orphan) allosteric sites.

In contrast to caspase-1 above, using a RRIN with cutoff radius of 6 Å, we find that the phosphorylation site of CheY is identified as a hot-spot: the average quantile score of allosteric residues is much higher for the rest of the residues 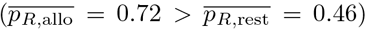. The RRIN detection is robust over a range of cut-off radii between 6Å-16Å (Table S6 and Fig. S5). This result suggests that sometimes (as for CheY) it is the *topology* of the protein structure that is important for signal propagation, whereas in other cases (as for caspase-1) the specific *atomistic structure* given by the *chemistry* of the side-chain interactions matters for allosteric propagation. Our all-atom methodology incorporates both aspects consistently.

**FIG. 2.**
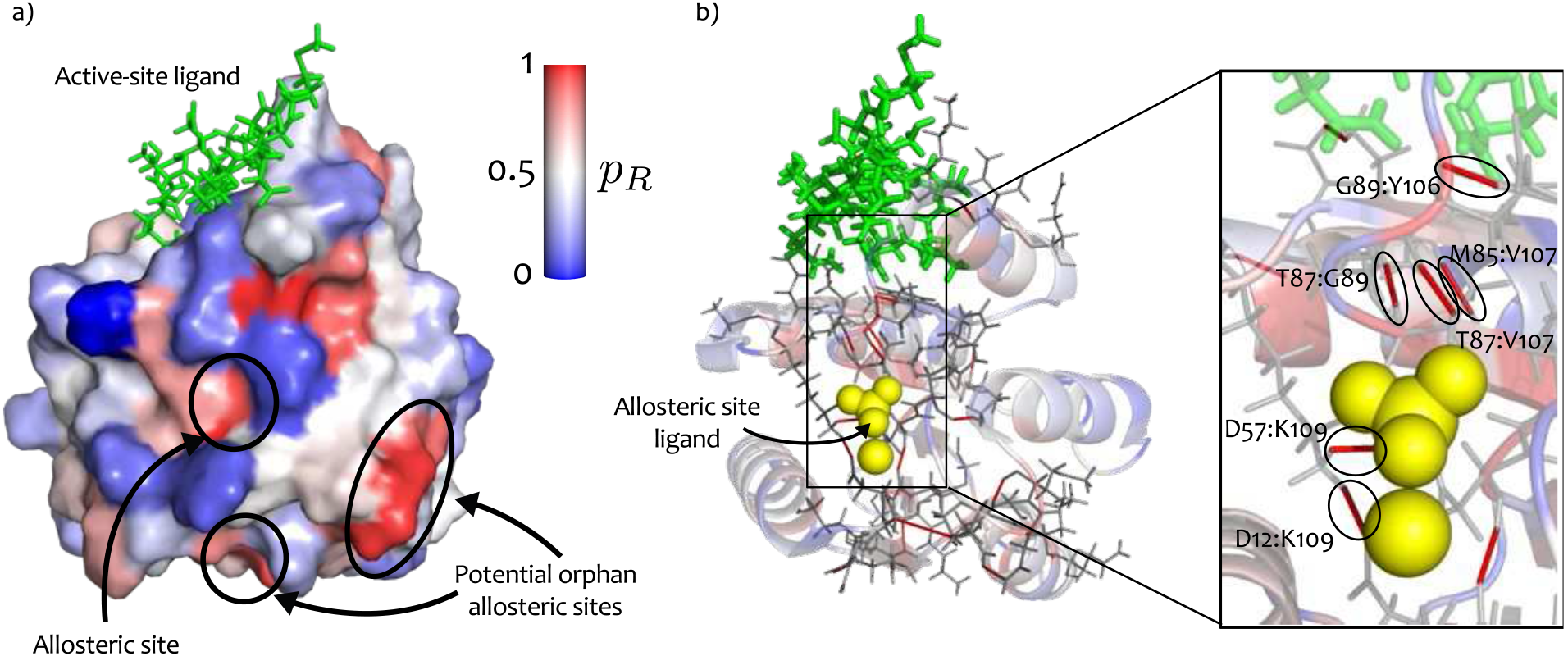
Allosteric phosphorylation site in CheY is identified by its high propensity. (a) Residue quantile scores *p_R_* are mapped onto the surface of CheY. The allosteric phosphorylation residue D57 is identified as a hot-spot. We identify two other distant sites, which could serve as potential orphan targets for allosteric effectors. (b) The top 3% of bonds by quantile score (i.e., *pb* ≥ 0.97) are indicated on the structure. The blow-up shows high-quantile score non-covalent bonds that form propagation pathways between the allosteric ligand (yellow spheres) and the ligand binding site (green).

**TABLE II.** Propensities of residues in CheY relative to the active site, ranked by quantile score (*p_R_* ≥ 0.90). Residues marked with a star are within 3.5 Å of the allosteric effector.

| Residue | $\Pi_R^{\text{act}}$ | $p_R$ |
| --- | --- | --- |
| D12 | 0.0076 | 1 |
| E89* | 0.0370 | 0.984 |
| N62 | 0.0017 | 0.984 |
| D57* | 0.0094 | 0.968 |
| K45 | 0.0015 | 0.968 |
| T87* | 0.0283 | 0.968 |
| M85 | 0.0321 | 0.968 |
| E35 | 0.0019 | 0.952 |
| L116 | 0.0189 | 0.952 |
| W58* | 0.0247 | 0.936 |
| L43 | 0.0030 | 0.921 |
| F124 | 0.0120 | 0.905 |
| L120 | 0.0189 | 0.905 |

#### 2. Comparing propensities of active and inactive structures helps identify allosteric communication networks

To get a more detailed picture of the pathways involved in allosteric communication, we examined the specific bonds with high propensity in the structure of fully activated CheY (1F4V). Considering high quantile scores (*p_b_* ≥ 0.97), we find several bonds connecting the allosteric phosphorylation site to the key binding site residue Y106 (Fig. 2b). One pathway comprises bonds between T87:E89 (*p_b_* = 0.991) and E89:Y106 (*p_b_* = 0.977), whereas a second pathway is formed by K109, which has high quantile score bonds with D12 (*p_b_* = 1) and D57 (*p_b_* = 0.993). These residues have been discussed extensively in the biochemical literature and are known to be crucial for allosteric signalling (see Discussion).

In addition to fully activated CheY, we also studied four additional structures corresponding to conformations of CheY across a range of activation stages (details of the PDB files and an in-depth comparison is given in SI Section 3). Importantly, the profiles of bond-to-bond propensities are similar across all conformations (Fig. S1), highlighting the robustness of the propensity scores to local dynamical rearrangements across different conformations. In particular, the propensities of residues in the active (1F4V) and inactive (3CHY) conformations show a strong positive correlation (*r* = 0.94, Fig. 3a). Using Cook’s distance, a well-known method for detecting influential points in linear regression [49], we identified E89, N94, T87, A98, and W58 as the residues with highly increased propensity in the active conformation as compared to the inactive conformation. Superposition of the active and inactive structures shows that the large displacement of E89 causes the formation of a tighter network of interactions involving N94, T87, and W58 in the active conformation (Fig. 3b). Interestingly, the propensity of the allosteric phosphorylation site D57 is similar in the active and inactive conformations; in the inactive conformation, D57 forms a stronger hydrogen bond with K109 than it does in the active conformation, yet the weakening of this bond in the active conformation is compensated for by the formation of the network involving W58 and E89. Hence activation induces a structural re-arrangement of the network of bonds that connect the phosphorylation site to the active site.

**FIG. 3.**
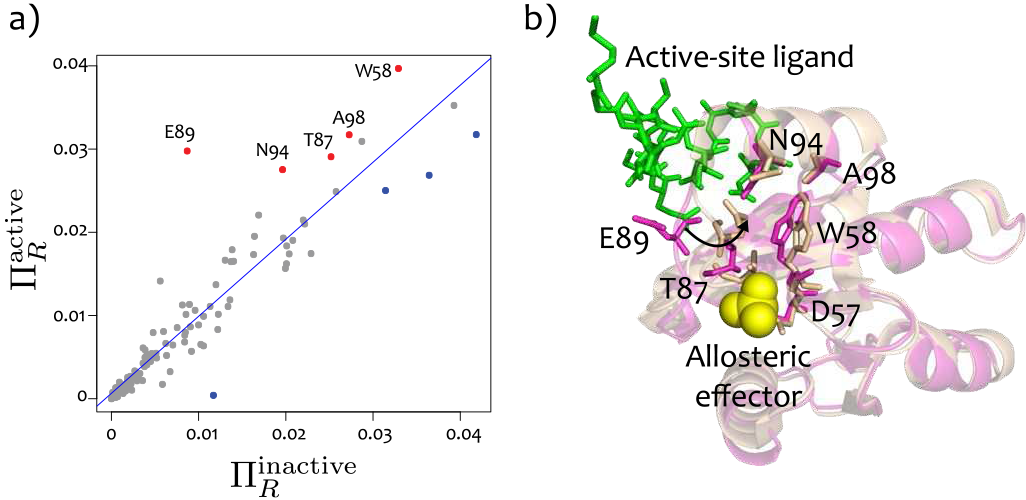
Comparison of residue propensities between active and inactive conformations of CheY. (a) The propensities most increased in the active X-ray structure (1F4V) as compared to to the inactive X-ray structure(3CHY), as identified by Cook’s distance, are coloured red and labelled. (b) Superposition of active (1F4V - beige) and inactive (3CHY - pink) conformations. The residues found in (a) form a pathway between the allosteric site and the ligand binding surface.

#### 3. Variability of bond-to-bond propensities in NMR ensembles uncovers transient effects in the allosteric network

CheY exists in dynamic equilibrium between its active and inactive conformations. Indeed, X-ray structures have revealed an intermediate conformation with only the binding site adopting the active conformation [50, 51].

To explore the effect of small structural changes on the propensities of residues of CheY, we analysed 20 NMR structures of the inactive conformation *apo*-CheY (PDB: 1CYE) and 27 NMR structures of the fully activated CheY bound to the phosphate mimic BeF_3_ (PDB: 1DJM). We calculated the average 〈Π_*R*_〉_NMR_ and the standard deviation SD(Π_*R*_)_NMR_ of the propensity of each residue over the ensemble of NMR structures. We then compared these properties computed over the NMR ensemble against those obtained from the X-ray structure.

The results of this comparison (NMR ensemble *vs*. X-ray structure) are different for the inactive and active structures, suggesting that the dynamical reconfigurations have a (consistent) effect on our measure. For the inactive *apo*-CheY, the average NMR propensity over the ensemble 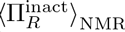 for each residue is strongly correlated (*r*^2^ = 0.96) with its X-ray propensity 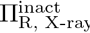 (Fig. S2a). For the active Che-Y, however, the correlation is weaker (*r*^2^ = 0.84, Fig. S2b). McDonald *et al* [52] have suggested that phosphorylation causes a slight increase in the flexibility of CheY, as signalled by increased B-factors and root mean square fluctuations (RMSF) across the NMR ensemble for active CheY. This enhanced flexibility may account for the greater difference between the NMR ensemble and the X-ray structures for the active conformation.

The variability of the propensity of each residue, computed from the NMR active ensemble, is shown in Fig. 4a. Among the residues with high (top 10%) NMR standard deviation 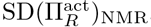, we find W58, T87, E89, and K109, which were also found to have high propensities in the active X-ray structure. These residues are known to be functionally relevant, and recent NMR relaxation-dispersion experiments have suggested that they form part of an allosteric network undergoing asynchronous local switching [52]. Other residues with high NMR standard deviation are A101, R73, L116, K119, and N121. Of these, A101 lies in the alpha-helix forming the top half of the ligand binding site, and the high variance of A101 and R73 can be explained by an unstable hydrogen bond between the two residues, which is transiently present across the active ensemble. On the other hand, L116 and N121 lie in the alpha-helix forming the other side of the FliM binding site: L116 forms a transient alpha-helical hydrogen bond with the ligand binding residue K119, and N121 forms fluctuating hydrogen bonds with residues in, and adjacent to, the active site (Fig. 4b).

**FIG. 4.**
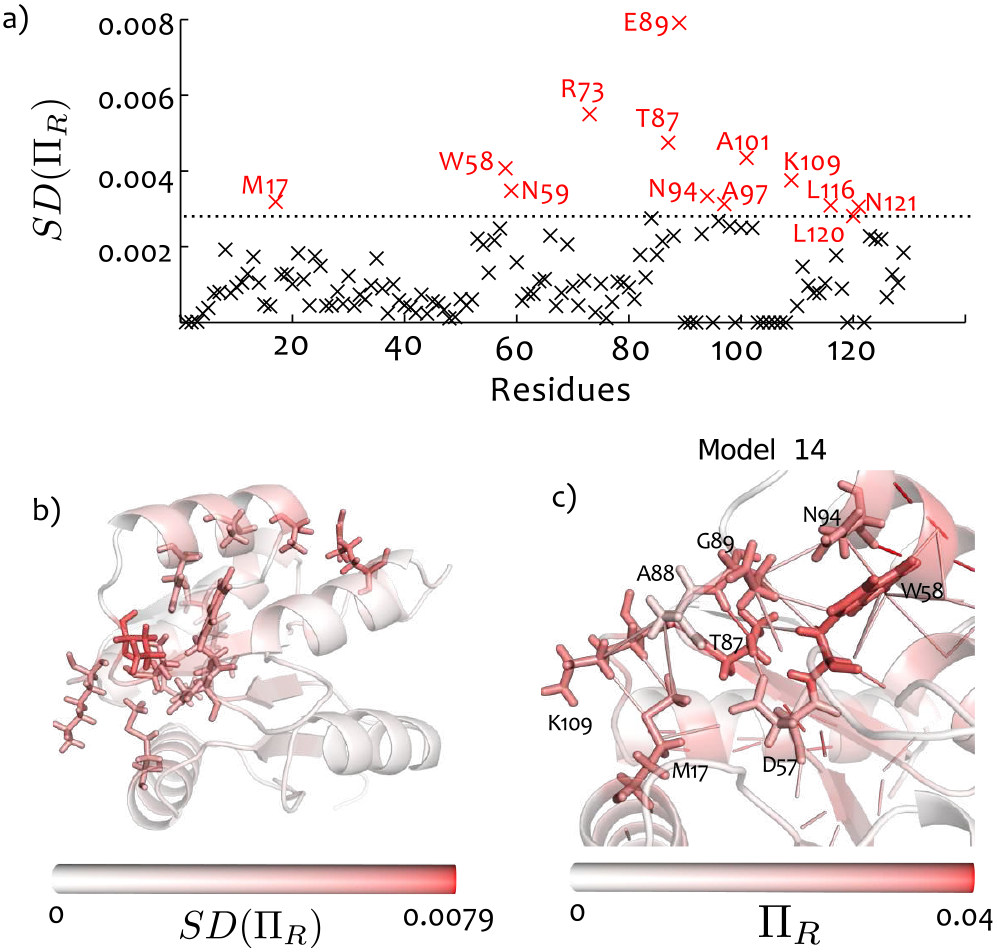
Increased variability of the propensity in NMR structures of active CheY reveals additional relevant residues. (a) Standard deviation of the residue propensities recorded over the NMR ensemble of 27 conformations corresponding to active CheY. The dashed line separates the top 10% of the residues by *SD*(Π_*R*_). Residue M17 has high NMR variability, although it was not identified in the X-ray structure as having high Π_*b*_. (b) The residues with high standard deviation are indicated on the structure, coloured by their NMR standard deviation. (c) Interactions coupling M17 to Y106 and the active site is shown in one of NMR conformations (model 14) of the active CheY. Residues coloured by their propensity Π_*R*_ in this particular conformation.

The large NMR variability of residue M17, which is 15Å away from the active site, is of particular interest. CheY is intolerant to mutation of M17 [53, 54], and it has been recently reported that this mutation causes chemical shift changes at Y106 [55], a key residue in the distant FliM binding site. Our analysis shows that the propensity of M17 is higher in the active structure (both NMR and X-ray) than in the inactive structure: 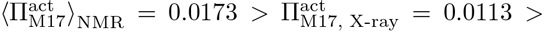 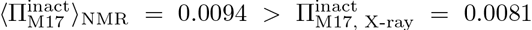. Furthermore, the NMR standard deviation of the propensity is higher in the active than in the inactive ensemble: 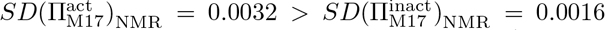. All these results indicate that phosphorylation (i.e., activation) causes transient pathways to form between M17 and the active site which are not observed in the X-ray structure. By examining bonds with high propensity between M17 and Y106, we visually uncover a communication pathway involving residue K109 and residues in the flexible *α*4 — *β*4 loop: T87, A88, and E89. Indeed, when we examine the individual NMR conformation in which M17 has the highest propensity, M17 bonds directly with A88 and is indirectly connected to T87 through a hydrogen bond with K109 (Fig. 4c). This suggests that M17 is transiently coupled to Y106 through a network of hydrogen bonds and hydrophobic contacts not captured in the active X-ray structure. In general, the transient making-and-breaking of particular bonds in the NMR ensemble translates into highly variable propensities associated with functionally important allosteric residues.

### C. Structural water molecules are crucial to the allosteric communication network in h-Ras

The enzyme h-Ras is a GTPase involved in signal transduction pertaining to cell-cycle regulation [56]. Crystallographic evidence shows that calcium acetate acts as an allosteric activator in this process [57]. By comparing the calcium acetate-bound structure to the inactive structure, Buhrman *et al* have proposed a network of hydrogen bonds, involving structural water molecules, linking the allosteric site to the catalytic residue Q61 [57].

**TABLE III.** Top bonds ranked by propensity quantile score for h-Ras (*p_b_* ≥ 0.99)

| Bond | $\Pi_b$ | Distance (Å) | $p_b$ |
| --- | --- | --- | --- |
| Q99:HOH727 | 0.0051 | 14.8 | 0.9991 |
| K117:G13 | 0.026 | 2.76 | 0.9983 |
| HOH727:S65 | 0.0067 | 12.2 | 0.9974 |
| R164:E49 | 0.0013 | 25.0 | 0.9974 |
| I21:S17 | 0.019 | 4.83 | 0.9965 |
| D47:R161 | 0.0015 | 21.6 | 0.9948 |
| H27:Q25 | 0.0075 | 10.8 | 0.9940 |
| V8:L56 | 0.0010 | 9.05 | 0.9940 |
| R161:D47 | 0.0013 | 21.6 | 0.9931 |
| I24:K42 | 0.0035 | 14.8 | 0.9922 |
| Q22:A146 | 0.017 | 5.09 | 0.9905 |

We have calculated the propensities and quantile scores of hRas (bound to substrate and allosteric activator, PDB code: 3K8Y) for two scenarios: with and without inclusion of structural water molecules in the graph. In the absence of water (Fig. 5a left), we find no bonds or residues with high quantile scores near the allosteric binding pocket. When we include the 8 molecules of structural water present in the PDB file, we identify a high quantile bond between the allosteric site residue Y137 and H94, and a pathway involving a structural water molecule that connects the allosteric region to a catalytic residue (Fig. 5b). In Table III, we show that the Q99-water and S65-water bonds involved in this pathway have 1st and 3rd highest quantile scores out of the 1159 weak interactions in the protein.

This water-mediated link between Q99 and S65 connects the allosteric binding pocket on helix 3 with the helical structure known as the switch 2 region, at the bottom of which lies Q61, which has been identified as a key catalytic residue [57]. Our results thus suggest that structural water plays a crucial role in coupling the allosteric effector to the catalytic residue Q61.

### D. Absolute bond propensities against a reference set from the SCOP protein database

The quantile regression scores *p_b_* in the previous sections identify bonds with high propensities as compared to other bonds which are at a similar distance from the active site *within the same protein*. To assess the *absolute* significance of bond propensities, we have assembled a reference set of 100 protein structures from the SCOP database [58] (see SI, Section 4), and calculated the propensities with respect to the active site of all 465,409 weak bonds in this reference set (Fig. 6a). Because the propensities are dependent on both the distance from the active site, *d*, and the total number of weak interactions in the protein, *E*, we apply quantile regression against both *d* and *E* (as given by Eq. (15) in Materials and Methods) to obtain fitted quantiles for the reference set. The quantiles computed from this reference set can then be used to obtain absolute bond propensity scores, denoted 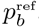, for any given protein without recomputing the regression.

We have obtained the absolute quantiles 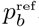 for the propensities of the three proteins (caspase-1, CheY, and h-Ras) studied above (Fig. 6b). Reassuringly, the significant bonds are also found to be important according to the absolute measure, with a strong correlation between quantile scores and absolute bond quantile scores (Fig. S3). Visualising the bonds with 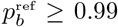 shows they form pathways between the active and allosteric sites (Fig. 6c). These results confirm that these bonds are important not only relative to other bonds and residues within each of the respective proteins, but also in absolute terms when compared to the protein reference set.

**FIG. 5.**
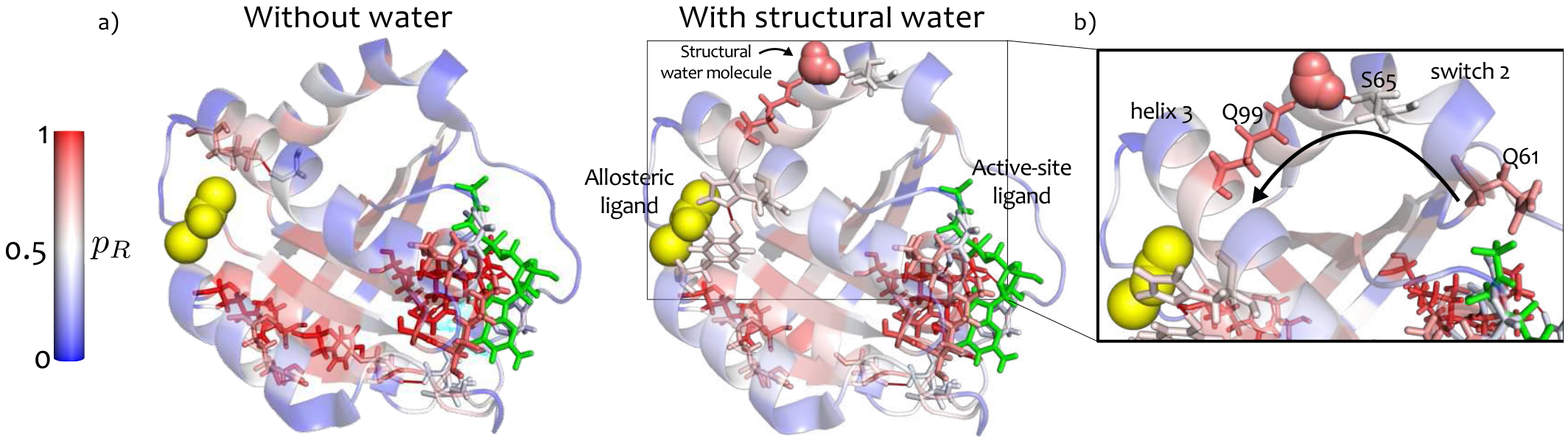
Structural water molecules are essential for the allosteric pathway in hRas. (a) Top percentile bonds by propensity quantile score (*p_b_* ≥ 0.99) are shown on the structure: the left panel shows pathways identified without the inclusion of water molecules, and the right panel when structural water molecules are included in the graph. The structural water allows the formation of a pathway between the bottom of the switch 2 region and the top of helix 3, where the allosteric binding site is situated. The crucial water molecule which connects Q99 and S65 is indicated. (b) Blow-up indicating details of the pathway formed by Q99, a water molecule and S65, linking the allosteric pocket to the switch 2 region. The catalytic residue Q61 is shown at the bottom of switch 2.

### E. Validating the propensity measure: predicting allosteric sites in an extended set of proteins

To test the validity of our methodology, we have computed the bond propensities for an additional 17 proteins known to exhibit allostery. Ten of these proteins were taken from a benchmark set collected by Daily *et al* [59] and a further 7 were obtained through an extensive literature search. (Five proteins in Ref. [59] could not be used either due to the presence of non-standard amino-acids, to the absence of an allosteric ligand, or to a mismatch between the oligomeric state of the active and inactive structures.) The details and structures of all 20 proteins analysed in the paper are given in the SI (Table S2 and Figure S4).

For each protein, we calculated the propensity quan-tile scores of all its bonds and residues, both intrinsic (*p_b_*, *p_R_*) and absolute 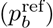, with respect to their active site. Again, no *a priori* knowledge about the allosteric site was used. Figure 7 shows the structures of the 20 proteins coloured according to the residue quantile score *p_R_*, with the allosteric sites marked with spheres. To validate our findings on this test set, we used the location of the allosteric site *a posteriori* and evaluated the significance of the computed allosteric quantile scores according to four statistical measures (Fig. 7a–d). See Section IV D for a full description and definitions.

All combined, the allosteric site is detected significantly by at least one of the four measures in 19 out of 20 proteins in the test set, and is detected by three or more of our measures in 15 out of 20 proteins in the test set. The full numerical values are given in the SI (Table S3). In practice, all statistical measures provide important and complementary information about the distribution of bond propensities, and can be used in conjunction for the detection of allosteric sites.

## III. DISCUSSION

Using a description of protein structural data in terms of an atomistic energy-weighted network with both covalent and non-covalent bonds, we have defined a graph-theoretic measure of bond propensity and used it to identify allosteric sites in proteins without prior information as to their location. Our propensity measure identifies bonds that are strongly coupled to the active site via communication pathways on the protein graph, even if they might be separated by large geometric distances. Allosteric sites correspond to hot spots, i.e., sites with high propensity to perturbations generated at the active site, as measured by their quantile score relative to other sites in the protein that are at a similar distance from the active site. This finding suggests that the structural features embedded in the architecture of the protein are exploited so as to enhance the propagation of perturbations over long distances.

By using a representative reference set of 100 proteins randomly assembled from the SCOP database, we also computed absolute quantile scores to further confirm the significance of bond propensities. One advantage of this absolute measure is that the quantile regression over the reference SCOP set does not need to be re-calculated, and the absolute bond quantile scores in any protein of interest can be obtained directly against them, thus reduces the analysis time even further.

We have validated our method against a test set of 20 allosteric proteins without using any *a priori* information of their allosteric sites. We used our propensity quantile scores and a structural bootstrap to define four statistical measures of significance based on the average and tail of the distribution of bond propensities in the allosteric site. The allosteric site is detected for 19/20 proteins, according to at least one statistical measure, and for 15/20, according to at least three of our four statistical measures. These findings indicate the robustness of the bond-to-bond propensity as a predictor of allosteric sites, which could be used to guide structure-based drug discovery efforts, e.g., by ranking potential binding sites based on their allosteric potential. Our method also uncovers hot spots not previously identified as allosteric sites (see our results for CheY in Fig. 2). Hardy and Wells have discussed the existence of ‘orphan’ or ‘serendipitou’ allosteric sites, i.e., sites targeted by as-yet undiscovered natural effectors or open for exploitation by novel small molecules [8]. The identified sites could thus provide targets for mutational analysis or allosteric small-molecule inhibition.

**FIG. 6.**
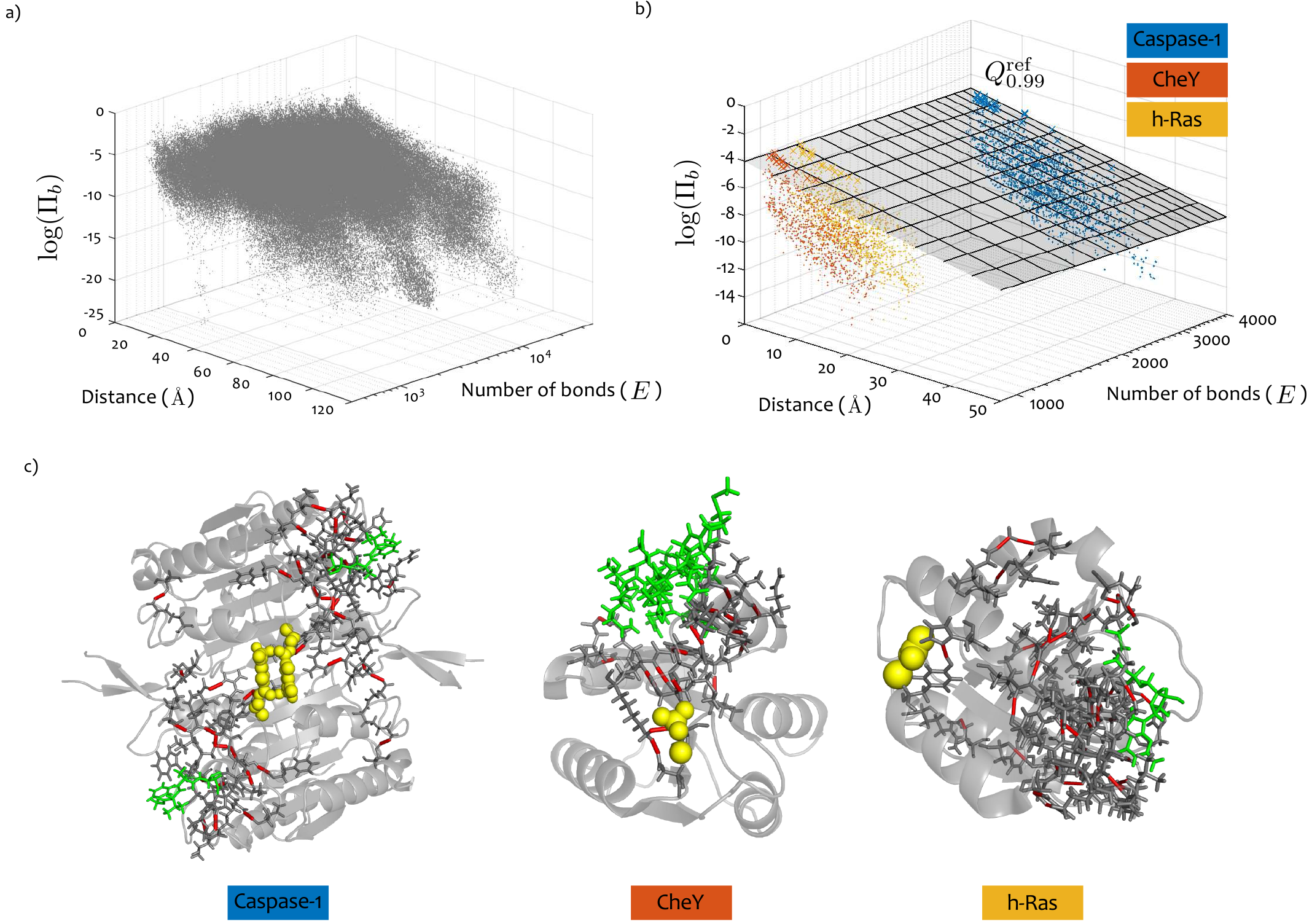
Absolute propensities: calibration against the SCOP reference set. (a) The logarithm of the bond propensity
log(Π_*b*_) of all 465,409 weak bonds in the reference set (100 proteins from the SCOP database) plotted against *d*, the distance from their corresponding active site, and *E*, where *E* is the number of weak bonds in the corresponding protein. (b) The log propensities log(Π_*b*_) for caspase-1 (blue), CheY (orange), and h-Ras (yellow) are plotted together with the plane defining the 99th quantile fit obtained by solving the optimisation Eq. (15) against the SCOP set of bonds shown in (a). For each of the three proteins, there are bonds lying above the 99th quantile plane. (c) The bonds above the plane in (b) have 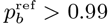 and are marked in red on the corresponding protein structures (active site ligand in green, allosteric ligand as yellow spheres). The bonds thus identified play key allosteric roles, in agreement with the *intrinsic* results in previous sections.

We have exemplified the use of atomistic propensities with the detailed analysis of three proteins (caspase-1, CheY, and h-Ras), focussing on the contribution of high propensity bonds to pathways (or networks) of weak bonds linking the active and allosteric sites. The weak bond network we found in caspase-(E390/R286/S332/S339/N337) has been previously tested experimentally and shown to be functionally important [48]. In CheY, we found that bonds between T87:E89 and E89:Y106, with very high quantile scores, are key to an important pathway for transmission of the signal induced by phosphorylation, also consistent with experimental evidence [50, 52, 60]. We also found a second pathway in CheY involving the bond K109:D57 (3rd highest quantile score). Interestingly, mutation of K109 abolishes chemotactic activity [53] and has been proposed to form part of the post-phosphorylation activation mechanism [61]. Our analysis of bond propensities across active/inactive conformations and NMR data further confirmed that K109 forms a central link in the communication between the phosphorylation and binding sites in CheY.

**FIG. 7.**
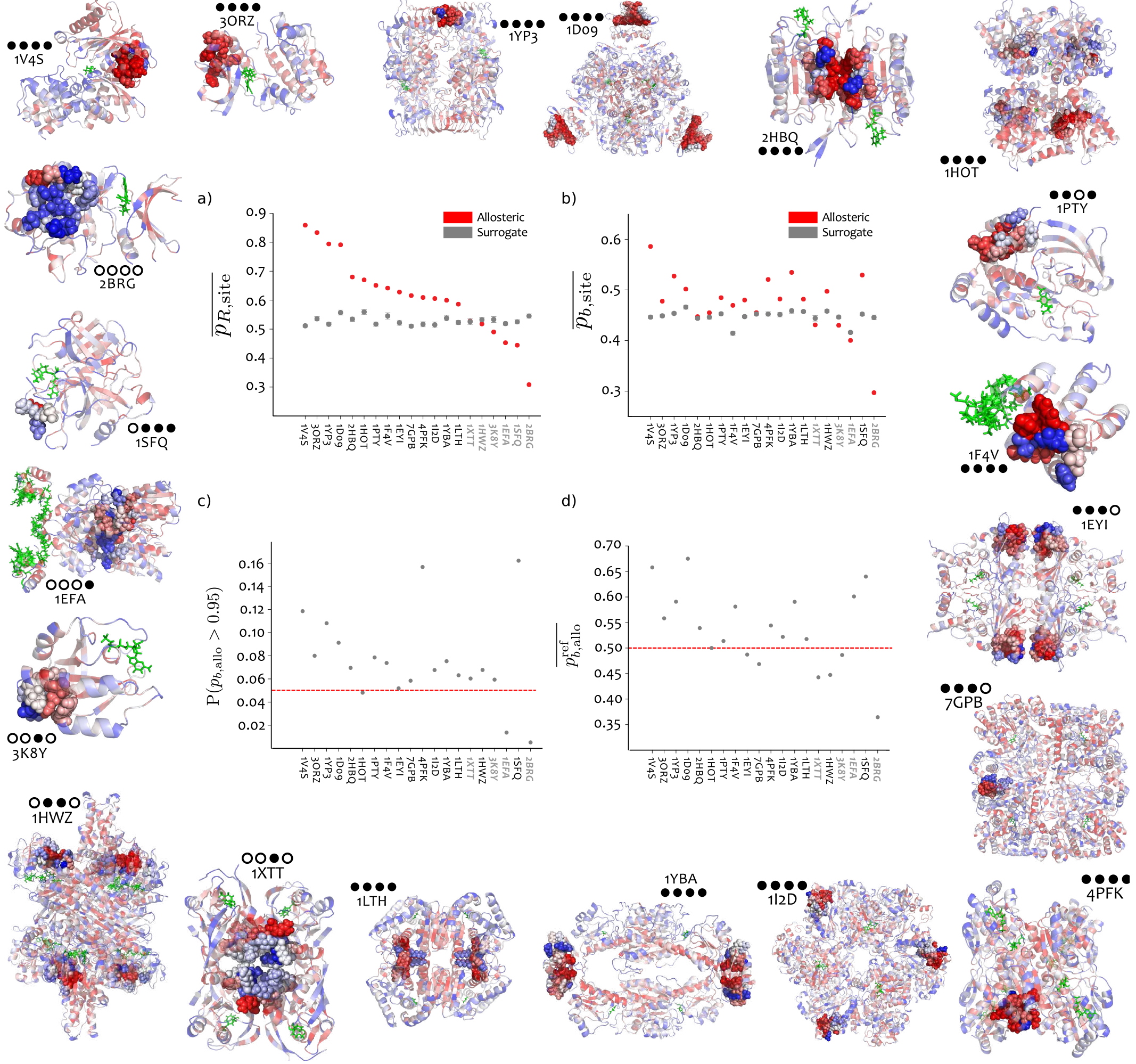
Prediction of allosteric sites based on bond-to-bond propensity for a test set of 20 allosteric proteins. The structures of the 20 proteins in the test set (labelled by PDB code) have their residues coloured by their quantile score *p_R_*, and the allosteric site is shown as spheres. For full details of these proteins, see Table S2 in the SI. The four statistics computed from our propensity are showed in the centre: (a) average *residue* quantile scores in the allosteric site 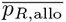 (red) compared to the average score of 1000 surrogate sites 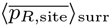 (grey), with a 95% confidence interval for the average from a bootstrap with 10000 resamples (see Section IV D 1); (b) average *bond* quantile scores in the allosteric site against the equivalent bootstrap of 1000 surrogate sites; (c) tail of the distribution of bond propensities, i.e., proportion of allosteric site bonds with quantile scores *p*_*b*,allo_ > 0.95. Proteins above the *expected* proportion of 0.05 (red line) have a larger than expected number of bonds with high quantile scores; (d) average *reference* bond quantile score in the allosteric site 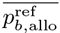. The red dotted line indicates the expected value of 0.5, and proteins above this line have a higher than expected reference quantile score. For the numerical values of all measures see Table S3 in the *SI*. The four circle code by each protein indicates whether the allosteric site is identified (filled circle) or not identified (open circle) according to each of the four measures (a)-(d). 19/20 allosteric sites are identified by at least one measure, and 15/20 sites are identified by at least three of four measures.

Determination of protein structures from NMR solution experiments results in multiple models, each consistent with experimentally-derived distance restraints. The resulting ‘ensemble’ of structures should be interpreted with caution, since variation could be due to actual flexibility and thermal motion during the experiment, or to inadequate (or under-constrained) interatomic distance restraints. Hence the set of NMR structures is not a true thermodynamic ensemble. However, our analysis suggests that the variation within the NMR structures can reveal functionally relevant information. For CheY, residues with highly variable propensities across the NMR ensemble (E89/W58/T87/E89/K109) coincide with those forming an asynchronously switching allosteric circuit after phosphorylation, as revealed by NMR relaxation-dispersion experiments [52]. We also identify residue M17 as having high propensity in the NMR ensemble due to the presence of a transient network of interactions. This may explain experiments showing that mutation of M17 has a functional effect and causes chemical shift changes at Y106 [55].

Comparing the results across conformations indicates that propensities are fairly robust to local dynamic fluctuations, as seen by the strong correlation between active and inactive conformations and across NMR structures (Fig. 3 and Figs. S1 and Fig. S2). As an additional confirmation of its robustness, we show in SI (Section 6, Tables S4 and S5) that the propensities, and the ensuing identification of significant residues and bonds, are generally robust to both randomness in the bond energies and to the breakage of a large proportion of weak interactions. On the other hand, as discussed above, our graph-theoretic analysis shows that further information about residues and bonds can be obtained by evaluating the highest variations induced by dynamical and structural variations. A fuller investigation of the effect of dynamics on the calculated propensities using experimental data (NMR, conformational studies) and complemented with the analysis of molecular dynamics simulations would thus be an interesting area for future research.

The role of structural water molecules in mediating allosteric communication has so far received limited attention. In a recent study of a PDZ domain, Buchli *et al.* suggest that changes in water structure could be responsible for mediating communication with remote parts of the protein [62]. Our analysis of h-Ras found that *structural* water molecules in the protein graph are necessary to reveal a pathway linking the allosteric and active sites. These results and the findings of Buchli *et al.* suggest that novel methods to study interaction networks between proteins and water are worth investigating. However, beyond including structural water when present in experimental structures (as in h-Ras here), the addition of bulk water would require the simulation of hydration, including energy minimisation and equilibration steps. This could constitute another direction of future research, since the computational efficiency of our method would make it possible to analyse all-atom representations of such hydrated structures.

To what extent does the identification of the allosteric site require an atomistic, chemically detailed construction of the graph? To answer this question, we applied our propensity measure to residue-residue interaction networks (RRINs), the coarse-grained residue-level models used in almost all previous network analyses of proteins. For caspase-1, we found that allosteric residues are not found significant in RRINs (across several different cut-off radii), whereas, on the other hand, the allosteric site of CheY was consistently detected in both the atomistic and residue-level descriptions. This indicates that both coarser topological features, as well as more detailed chemical communication pathways can be relevant depending on the protein; e.g., in caspase-1, the binding of the allosteric ligand perturbs a network of strong hydrogen bonds and salt-bridges as identified in our analysis. Therefore, the atomistic graph with detailed physicochemical information can in some cases provide important features underpinning the communication features of the protein. The analysis of coarse-grained models with a variety of cut-off radii for all 20 proteins in our allosteric test set in SI Section 7 confirm that the outcome for RRINs varies for each protein and can also be dependent on the choice of cut-off radii [38]. We would like to emphasise, however, that our propensity measure is principally agnostic to the protein network model under analysis, thus allowing for the evaluation of distinct graph-construction techniques (e.g., atomistic vs coarsegrained) or the use of different force-fields. Again, this would open another interesting avenue for future work.

Finally, it is important to remark that our method is computationally efficient. To obtain the bond-to-bond propensities, we only need to solve a sparse linear system (Eq. 6) involving the (weighted) Laplacian of the protein graph. As discussed in Section IV A 3, recent algorithmic advances allow us to solve such linear systems in *almost linear time* [42, 43]. Hence protein complexes of ~ 100,000 atoms can be run in minutes on a standard desktop computer. We can thus maintain atomistic detail, yet analyse large biomolecular complexes that are intractable for traditional computational methods.

## IV. MATERIALS AND METHODS

### A. Mathematical derivation of the bond-to-bond propensity

#### 1. Fluctuations and the edge-to-edge transfer matrix of a graph

The edge-to-edge transfer matrix *M* was introduced in Ref. [41] as a non-local edge-coupling matrix for the analysis of weighted undirected graphs, based on the concept of flow redistribution. In that work, it was shown that the element *M_ij_* reflects the effect that an injected flux on edge *i* has on the flux along edge *j* after the fluxes are redistributed over the whole graph when at equilibrium. Alternatively, *M* can be understood as a discrete Green’s function in the *edge space* of the graph. See [41] for detailed derivations and applications.

In this paper, we derive a complementary interpretation of the matrix *M*. As shown below, the edge-to-edge transfer matrix can be understood as describing how the fluctuations of the edge weights propagate through the graph. This new re-interpretation underpins the work in this paper, as it highlights the importance of *M* for the analysis of bond fluctuations in biomolecules.

As our starting point, consider the well-known Langevin equation, sometimes denoted the *heat kernel equation* [63, 64]:

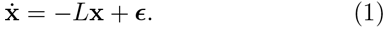

Formally, Eq. (1) has the same structure as the canonical model for scalar vibrations with nearest neighbour interactions encoded by the matrix *L* [29, 30]. Alternatively, Eq. (1) may be considered as a model of a diffusing particle transitioning like a random walker on the underlying graph structure represented by *L*. In contrast to residue level methods [33], the variable x is associated with *atomic* fluctuations, i.e., our graph model reflects an atomic description that incorporates physico-chemical interactions derived from the three dimensional structure of the protein recorded in a PDB file. The resulting graph contains energy-weighted interactions representing bonds in the protein, including both covalent bonds and weak interactions such as hydrogen bonds, salt bridges, hydrophobic tethers, and electrostatic interactions. For details of the graph construction see Section IV E and SI.

The matrix *L* is the graph Laplacian [39, 65]:

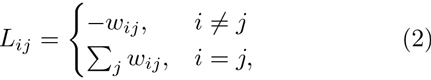

where *w_ij_* is the weight of the edge between nodes (atoms) *i, j*. In this case, *w_ij_* is the energy of the bond between both atoms. Thermal background fluctuations are modelled by ɛ, a zero mean white Gaussian noise input vector, i.e., a (simple) heat bath acting independently on all atomic sites with covariance matrix

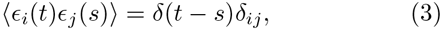

where *δ* stands for the Dirac delta function.

Instead of focusing on the *atomic* (node) variables x, we wish to study the coupling between *bonds*, and thus concentrate on the bond (edge) variables of the graph:

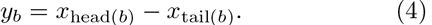

Clearly, *y_b_* describes the difference of the node variables at the endpoints of the associated bond *b*, i.e., a *fluctuation* associated with the bond between two atoms. The vector of bond fluctuations can be compactly represented in vector notation as:

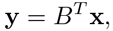

where *B* is the incidence matrix of the graph relating each edge variable to its corresponding node variables, i.e., *B_bi_* = 1 if node *i* is the head of bond *b*; *B_bi_* = —1 if node *i* is the tail of bond *b*; and *B_bi_* = 0 otherwise.

We can now calculate the cross-correlations between edge fluctuations as:

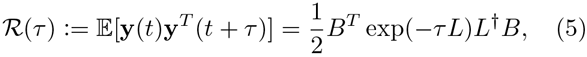

where *L*^†^ is the (Moore-Penrose) pseudoinverse of the Laplacian matrix. Each entry [ℛ(*τ*)]_*b*1*b*2_ describes how a fluctuation at bond *b*_2_ is correlated with a fluctuation at bond *b_1_* at time *τ*. See SI for a full derivation of Eq. (5).

Biophysically, we are ultimately interested in the *energy fluctuations* induced by bonds on other bonds. Therefore, we multiply the correlation matrix ℛ(*τ*) by the diagonal matrix of bond energies, *G* = diag(*w_b_*):

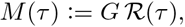

to obtain the matrix of *bond-to-bond energy correlations* with delay *τ*. Our measure of bond-to-bond propensity is obtained from the *instantaneous* correlations (i.e., *τ* = 0) leading to the edge-to-edge transfer matrix:

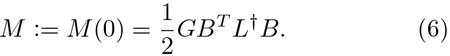

Note that the diagonal entries of *M* are indeed related to the average energy stored in the bond fluctuations: 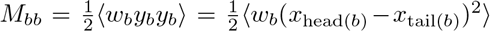. Likewise, the off-diagonal entries 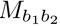 reflect how a perturbation at bond *b*_2_ affects another bond *b*_1_ weighted by the strength of bond *b*_1_. Hence the influence on a stronger bond is considered to be more important. Although we have not considered here time-delayed correlations (i.e., as a function of *τ*), this is an interesting direction for future research.

#### 2. Definition of the bond-to-bond propensity

To construct our measure of propensity, we only assume knowledge of the active site and proceed as follows. Let us consider all the ligand-protein interactions formed at the active site and compute their combined effect on each bond *b* outside of the active site:

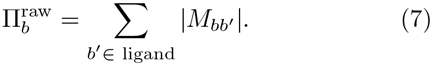

This raw propensity reflects how closely the active-site is coupled to each individual bond. Note that the computations include *all* the bonds in the protein (covalent and non-covalent). However, in the paper we only report the effect on weak bonds, as it is changes in weak-bonding patterns that usually drive allosteric response in proteins. Since different proteins have different numbers of bonds, we make the measure consistent by normalising the score:

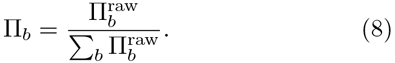

Throughout the manuscript, the quantity Π_*b*_ is referred to as the *propensity* of bond *b*; a measure of how much edge *b* is affected by the interactions at the active site. The propensity of a *residue* is defined as the sum of the (normalised) propensities of its bonds:

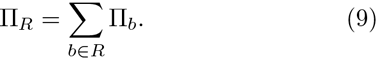

#### 3. Computational cost of bond-to-bond propensity

The computation of the propensities is efficient. Note that Eq. (8) requires the summation over columns of the *M* matrix corresponding to protein-ligand interactions. Crucially, we do not need to compute the full pseudo-inverse *L*^†^ in Eq. (6); we can instead solve a sparse linear system involving the graph Laplacian. Recent algorithmic developments [42, 43] have made this possible in almost linear time, 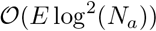, where *E* is the number of bonds (edges) and *N*_a_ is the number of atoms (nodes). Our method therefore is scalable to extremely large systems. Using the Combinatorial Multigrid toolbox written by Y. Koutis [66] (available at http://www.cs.cmu.edu/~jkoutis/cmg.html) propensities for all the bonds in proteins with ~ 100,000 atoms can be run in minutes on a standard desktop computer.

### B. Significance of propensities through quantile scores

To identify bonds (and residues) with high propensities relative to others at a similar distance from the active site, we use *quantile regression* [44], a technique of wide use in econometrics, ecology, and medical statistics. In contrast to standard least squares regression, which focusses on estimating a model for the conditional mean of the samples, quantile regression (QR) provides a method to estimate models for conditional quantile functions. This is important for two reasons: (i) the conditional distributions of propensities are highly non-normal; and (ii) we are interested not in the ‘average’ bond, but in those bonds with particularly high propensities lying in the tails of the distribution. Once the fitted models are obtained, the quantile score of a bond *p_b_* is a measure of how high the propensity Π_*b*_ is relative to other bonds in the sample which are at a similar distance from the active site.

Although QR goes back more than 200 years, it has only become widely used recently, due to the availability of computational resources. The mathematical basis of the method stems from the fact the *p*^th^ quantile, *Q_p_*, of a distribution is given by the solution of the following optimisation problem: given a sample 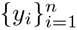 parametrically dependent on *m* variables x_*i*_ ∈ ℝ*^m^* with parameters *β*, the estimate of the conditional *p*^th^ quantile of the sample distribution is obtained by solving

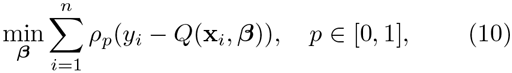

where *ρ_p_*(․) is titled absolute value function

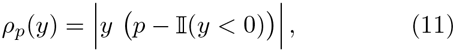

and 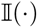 is the indicator function. If the dependence is assumed to be linear, *Q*(x_*i*_, *β*) = *β*_0_ + *β^T^*x_*i*_, the optimisation can be formulated as a linear program and solved efficiently through the simplex method to obtain 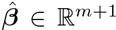, the estimated parameters defining the model [44].

In Sections IIA–IIC, we have applied QR to the propensities Π_*b*_ of bonds within each protein so as to take into account their dependence with respect to *d_b_*, the minimum distance between bond *b* and any bond in the active site:

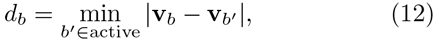

where the vector **v**_*b*_ contains the coordinates of the midpoint of bond *b*. Based on the observed exponential decay of Π with *d*, we adopt a linear model for the logarithm of the propensities and estimate the conditional quantile functions by solving the minimisation problem

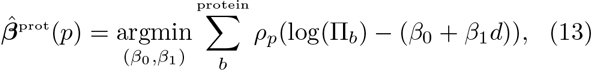

where the sum runs over the weak bonds of the corresponding protein. From the estimated model for the protein, we then calculate the quantile score of bond *b* at distance *d_b_* from the active site and with propensity Π_*b*_, by finding the quantile *p_b_* such that

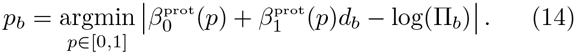

Similarly, in Section IID, we use QR to obtain *absolute* quantile scores of bonds and residues with respect to a reference set of 100 proteins from the SCOP database. In this case, the propensities are regressed against both the distance to the active site *d*, and the number of non-covalent bonds in the protein, *E*. Since the mean propensity scales as *E*^−1^, we also assume a power-law dependency of the quantiles. Hence, we solve

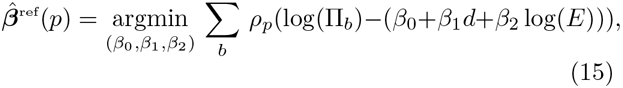

where the sum runs over all the weak bonds of all the proteins in the SCOP reference set. For each quantile *p*, the model is defined by the equation of a plane 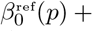 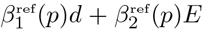 (Fig. 6b). The global quantile score 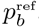 for bond *b* at a distance *d_b_* from the active site in a protein with *E_b_* non-covalent bonds is found by solving

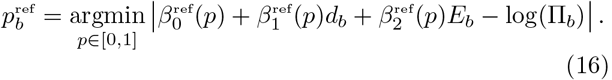

Quantile scores for residues are obtained by applying the same process to the propensities Π_*R*_.

The QR computations have been carried out using the R toolbox *quantreg* (http://cran.r-project.org/web/packages/quantreg/index.html) developed by R. Koenker [67].

### C. The SCOP reference set of generic proteins

The Structural Classification of Proteins (SCOP) database is a manually curated database which uses a hierarchical classification scheme collecting protein domains into structurally similar groups [58]. The major classes of cytoplasmic proteins in the database are *α, β,α/β, α+β*, and multidomain, covering all the major fold-types for cytosolic proteins. To obtain a representative set of proteins from the database, we randomly selected 20 proteins from each of the five classes. Note that we only include proteins for which there is a structure with a ligand bound to the active site. Our reference set thus covers a broad region of protein structure space. A table containing details of the 100 proteins selected can be found in the electronic *SI*.

For each protein in the dataset, we compute the distance from the active site, *d_b_*, and we calculate the propensity, Π_*b*_, for all its *E* weak bonds. Across the 100 proteins, we obtain a total of 465409 (*d, E*, Π_*b*_) 3-tuples corresponding to all the weak bonds in the proteins of the reference set (Fig. 6a). We then use QR to fit quantiles to this reference set, as given by Eq. (15). Note that the estimated quantile models, which are conditional on *d* and *E*, are now referred to the whole SCOP reference set and are not specific to any one particular protein. We then use the quantiles of the reference set to compare the bond propensities of any protein of interest and compute the *absolute* quantile score 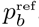 for each bond, as given by Eq. (16). This score measures how high the bond propensity is, given its distance from the active site and the number of weak bonds in the protein of interest, as compared to all the bonds contained in the wide range of proteins represented in the SCOP reference set.

### D. Statistical evaluation of allosteric site quantile scores

To validate our findings on the allosteric protein test set, we evaluated the significance of the computed quan-tile scores according to four statistical measures, based on the following metrics:

i. The average bond quantile score:

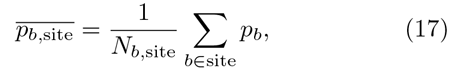

where *N_b,site_* is the number of bonds in the site.
ii. The average residue quantile score:

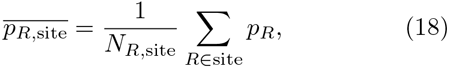

where *N_R,site_* is the number of bonds in the site.
iii. The proportion of allosteric bonds with *p_b_* > 0.95, denoted P(*p_b,allo_* > 0.95). Since the quantile scores are uniformly distributed, 0.05 is the expected proportion of bonds with quantile scores above 0.95.
iv. The average reference bond quantile score:

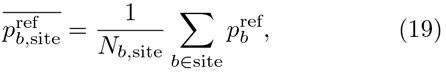

where *N_b,site_* is the number of bonds in the site.

These four measures are introduced to check robustly for the significance of the bonds in the allosteric site from distinct perspectives. If the functional coupling between active and allosteric sites is due to a cumulative effect of the entire allosteric site, then average quantile scores over all bonds in the allosteric site should be an accurate measure of its allosteric propensity. Measures (i), (ii) and (iv) capture this property at the level of bonds and residues for both intrinsic and absolute propensities. It is also possible that functional coupling to the active site is concentrated on a small number of high quantile score bonds, with most others only being involved in structural or energetic aspects of binding to the allosteric ligand and having low quantile scores. Our metric (iii), which measures the number of high quantile score bonds in the site, can capture this behaviour based on the tail of the distribution. Reassuringly, the four measures provide complementary, yet largely consistent outcomes.

#### 1. Structural bootstrapping

To establish the significance of the average quantile scores 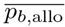 and 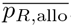, we assess them against random surrogate sites sampled from the same protein, used as a structural bootstrap. The surrogate sites generated satisfy two structural constraints: 1) they have the same number of residues as the allosteric site; 2) their diameter (i.e., the maximum distance between any two atoms in the site) is not larger than that of the allosteric site. The algorithm for generating these sites is described in Section S5 of the *SI*. For each protein, we generate 1000 surrogate sites and calculate their quantile scores 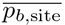 and 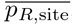. The average scores over the ensemble of 1000 surrogate sites 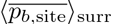 and 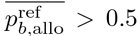, where the angle brackets denote the ensemble average, are then compared against the average residue quantile score of the allosteric site (Figure 7a, b). A bootstrap with 10000 resamples with replacement [68] was used to obtain 95% confidence intervals providing statistical signficance.

#### 2. Validation on the allosteric test set

Figure 7 (a)–(d) reports these four statistical measures for all 20 proteins analysed (see SI, Table S3 for the corresponding numerical data). Our results indicate robust identification of the allosteric sites in the test set. The quantile score of the allosteric site is higher than that of the surrogate sites and above the 95% bootstrapped confidence interval in 14 out of 20 proteins for the residue score, 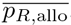, and for 16 out of 20 proteins for the bond score, 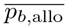 (Figure 7a, b). The proteins identified by both measures are almost coincident, with few differences: Glutamate DH (1HWZ) is significant according to the bond score and marginally below significance according to the residue score, whereas the opposite applies to Thrombin (1SFQ). The reason for these differences lies with the distribution of bond scores: in some cases, allosteric sites have only a few bonds with high quantile scores and many other less important bonds. When considered at the level of residues, this can lead to high *p_R_* scores; yet when bonds are considered individually through their *p_b_* scores, the high quantile scores are averaged out over the whole allosteric site.

To evaluate the presence of high scoring bonds, we compute the proportion of bonds with high quantile score P(*p_b,allo_* > 0.95) in the allosteric site, as compared to the expected proportion (0.05) above this quantile. The proportion of high quantile score bonds in the allosteric site is greater than expected in 17 of the 20 proteins (Fig. 7c). Of these 17 proteins, 16 coincide with those identified using the average scores reported above, and we additionally identify h-Ras (3K8Y). This finding confirms that allosteric sites consistently exhibit a larger than expected number of bonds with a strong coupling to the active site.

Finally, we compute the average *absolute* quantile score of the allosteric site 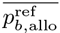 against the SCOP reference set (Figure 7d). The results are largely consistent with the intrinsic measure 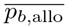: in 14/20 proteins, the absolute quantile score is greater than the expected 0.5, i.e. 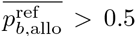. Yet some proteins (e.g., glutamate dehy-rogenase (1HWZ), fructose 1,6-bisphosphatase (1EYI), and glycogen phosphorylase (7GPB)) have high intrinsic quantile scores, as compared to other bonds in the same protein, but do not score highly in absolute value, as compared to the reference SCOP ensemble. This result highlights the fact that a site need not have a high absolute propensity, as long as its propensity is high in comparison with the rest of the protein it belongs to, so that the ‘signal’ from the site outweighs the ‘noise’ from the rest of the protein. Interestingly, the lac repressor (1EFA) has an allosteric site with large absolute propensity 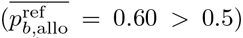 but non-significant intrinsic propensity.

### E. Construction of the atomistic graph

An in-depth discussion of the construction of the graph can be found in Refs. [39, 40], and further details are given in the SI, Section 2. Briefly, we use an atomistic graph representation of a protein, where each node corresponds to an atom and the edges represent both covalent and non-covalent interactions, weighted by bond energies derived from detailed atomic potentials. The covalent bond energies are taken from standard bond dissociation energy tables. Non-covalent interactions include hydrogen bonds, salt bridges, hydrophobic tethers and electrostatic interactions. Hydrogen bond energies are obtained from the DREIDING force-field [69]. Attractive hydrophobic interaction energies are defined between carbon and sulfur atoms, according to a hydrophobic potential of mean force introduced by Lin *et al* [70]. Electrostatic interactions with coordination ions and ligands are identified from the LINK entries in the PDB file, with bond energies assigned using a Coulomb potential.

To compare the results between our atomistic model and residue-level RRINs [33], we use coarse-grained network models obtained from the oGNM server [71]. A detailed comparison of results obtained with atomistic networks and RRINs is given in the SI Section 7.

We note that the main methodology (i.e., the propensity measure and methods developed in Sections IV A–IVB) is independent of the construction of the graph. Users are free to construct the network using alternative potentials (e.g., AMBER [72] or CHARMM [73]) or using coarse-grained networks.

## V. ACKNOWLEDGMENTS

BRCA was supported by a studentship of the EPSRC Centre for Doctoral Training under the Institute of Chemical Biology, Imperial College London. SNY and MB acknowledge support through EPSRC grant EP/I017267/1. We thank Keith Willison for suggesting h-Ras as an example and for helpful discussions.

## VI. AUTHOR CONTRIBUTIONS

BRCA, SNY and MB conceived the study. BRCA performed the numerical analysis and created the Figures. SNY and MB supervised the study. All authors contributed to developing the theoretical tools. All authors wrote and reviewed the manuscript.

## Supplementary Information

### Section 1 — Derivation of the graph-theoretical formula for edge fluctuations

We now derive in more detail Eq.(5), presented in Materials and Methods (Section IVA) in the main text. Let us consider the Langevin equation, Eq.(1) in the main text:

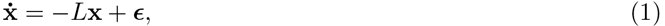

where ε is white Gaussian noise. Without loss of generality, we may assume that the system started initially from a condition x(—∞) = 0. A standard result from linear system theory is that the solution of equation (1) is given by:

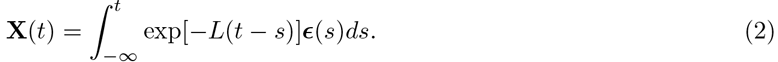

Since our input ε is random, X(*t*) is a random process, which we indicate by the upper-case notation. Likewise, the edge variables will be described by the random process:

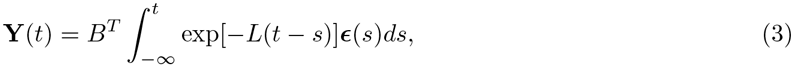

where *B* is the incidence matrix of the graph of the protein.

The autocorrelation of the process Y(*t*) for *τ* > 0 is then

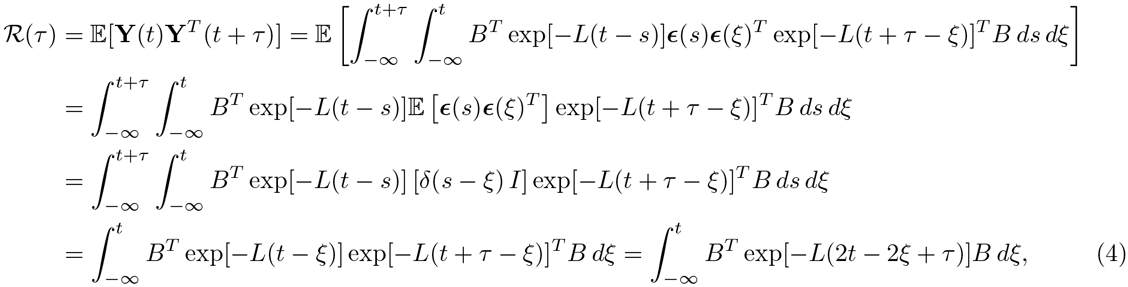

where we have used the fact that the noise vector ε is delta-correlated in time and across nodes (i.e., *I* = *δ_ij_* is the identity matrix). The last equality follows from fact that *L* = *L^T^*; hence it commutes and this implies that exp(*Lt*) exp(*Lt*)^*T*^ = exp(2*Lt*).

This integral can be computed using the eigendecomposition of the matrix exponential as follows:

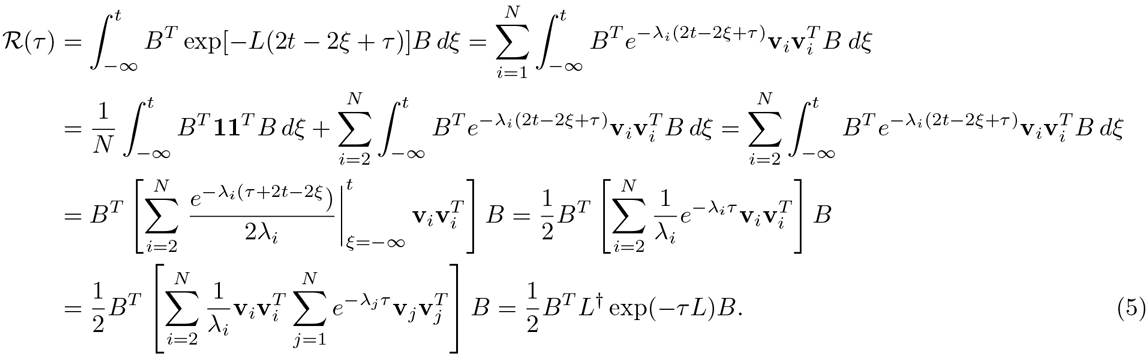

Here we have used the fact that the leading eigenvector of *L* associated with λ_1_ = 0 is the vector of ones (v_1_ = 1), which is in the null space of *B^T^*, i.e., *B^T^*1 = 0. In the last two equations we have made use of the orthonormality of the eigenvectors 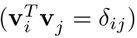, which implies that 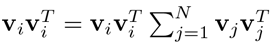.

### Section 2 — Construction of the atomistic protein network

As discussed in Materials and methods (Section IVE), the protein network is constructed by assigning edges between atoms which interact covalently and non-covalently. Each edge is weighted by the strength of the interaction. Covalent bond strengths are obtained from tables assuming standard bond lengths. We include three types of non-covalent interactions: hydrophobic interactions, hydrogen bonds, and electrostatic interactions. The assignment of bonds in the graph follows from the well established FIRST framework [1, 2]. More in detail:

- **Covalent bonds**: Covalent bonds are weighted according to standard bond dissociation energies given in Ref. [3].
- **Hydrophobic tethers**: Hydrophobic tethers are assigned between C-C or C-S pairs based on proximity: two atoms have a hydrophobic tether if their Van der Waals’ radii are within 2 Å. The hydrophobic tethers are identified using FIRST [4], which does not assign them an energy, and the energy is then determined based on the double-well potential of mean force introduced by Lin *et al* [5], which gives an energy of ≈ -0.8kcal/mol for atoms within 2 Å.
- **Hydrogen bonds**: The energies of hydrogen bonds were calculated using the same formula used by the program FIRST [4] and is based on the potential introduced by Mayo *et al* [6].
- **Electrostatic interactions**: Important electrostatic interactions between ions and ligands, as defined in the LINK entries of the PDB file, are added with energies derived from a Coulomb potential

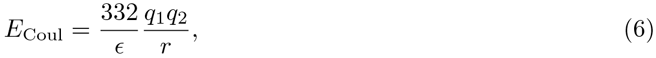

where *q*_1_ and *q*_2_ are the atom charges, *r* is the distance between them, and ε is the dielectric constant, which is set to ε = 4 as in Ref. [7]. Atom charges for standard residues are obtained from the OPLS-AA force field [8], whereas charges for ligands and non-standard residues are found using the PRODRG web-server [9].

An extended discussion of the construction of the atomistic graph can be found in Refs. [10, 11, 12]

### Section 3 — Propensities of CheY conformations: different activation states and NMR ensemble

In the following Table and Figures we give additional information about the active and inactive conformations and NMR data of CheY used in Section IIB of the main text.

#### 3.1 Active and inactive conformations of CheY

We calculated the propensities of residues for several CheY structures representing different activation states. Details of the different structures are given in Table S1 and a comparison of the perturbation propensities across the different structures is shown in Figure S1. As discussed in Section IIB.2, the propensities of the residues are strongly correlated across states. In the main text (Section IIB.2 and Figure 3), we concentrate on the comparison of 1F4V (active) against 3CHY (inactive).

**Table S1.**
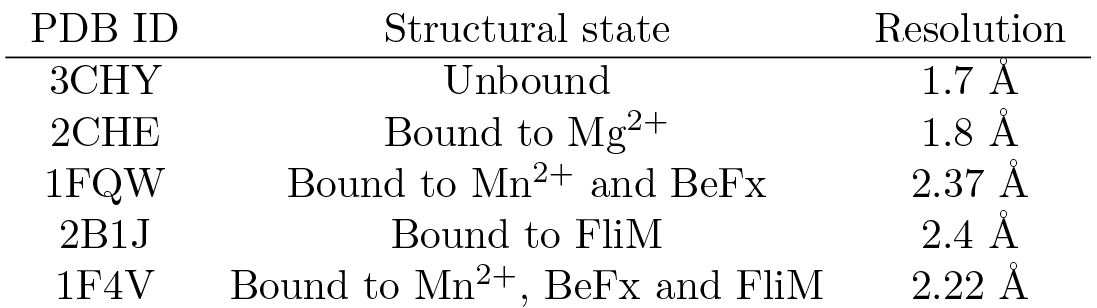
Details of X-ray structures of CheY analysed. The conformations correspond to different stages of activation.

| PDB ID | Structural state | Resolution |
| --- | --- | --- |
| 3CHY | Unbound | 1.7 Å |
| 2CHE | Bound to $\text{Mg}^{2+}$ | 1.8 Å |
| 1FQW | Bound to $\text{Mn}^{2+}$ and BeFx | 2.37 Å |
| 2B1J | Bound to FliM | 2.4 Å |
| 1F4V | Bound to $\text{Mn}^{2+}$, BeFx and FliM | 2.22 Å |

**Figure S1:**
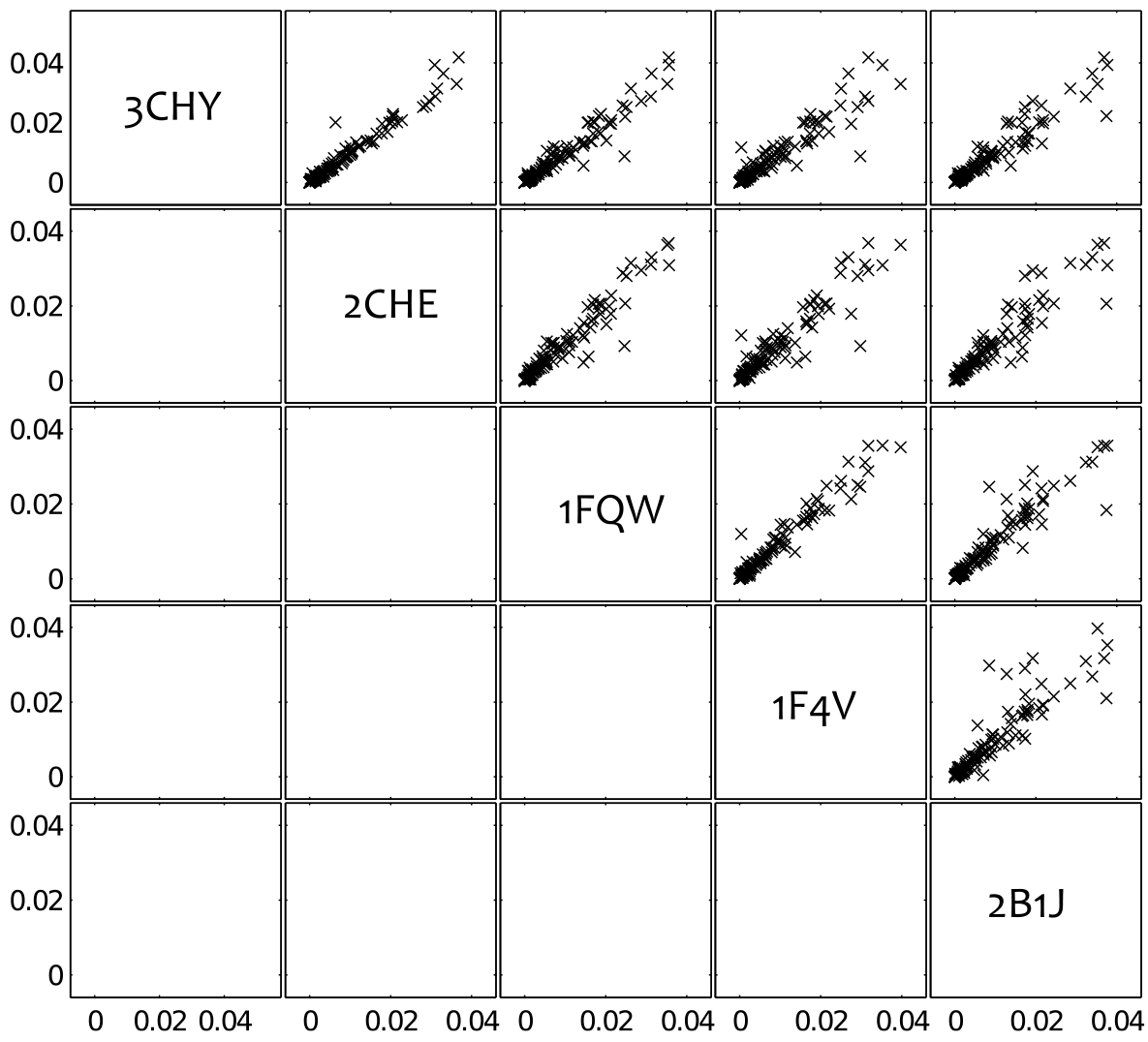
Propensities in different conformations of CheY. Comparison of propensities of residues in across different structures of CheY: unbound (3CHY); bound to Mg^2+^ (2CHE); bound to Mn^2+^ and phosphate mimic BeFx (1FQW); bound to Mn^2+^, BeFx and FliM (1F4V); and bound only to FliM (2B1J). The propensities of the residues are strongly correlated across states.

#### 3.2 CheY structures from NMR experiments

We also calculated the perturbation propensities of residues across two ensembles of NMR structures for active CheY (PDB ID: 1DJM; 27 structures) and inactive CheY (PDB ID: 1CYE; 20 structures). A comparison of the average propensity of each residue (averaged across the NMR ensemble) versus its propensity in the X-ray structure is shown in Figure S2 for both the active ensemble (1DJM) and the inactive ensemble (1CYE). This data is discussed in the main text (Section IIB.3) and summarised in Figure 4.

**Figure S2:**
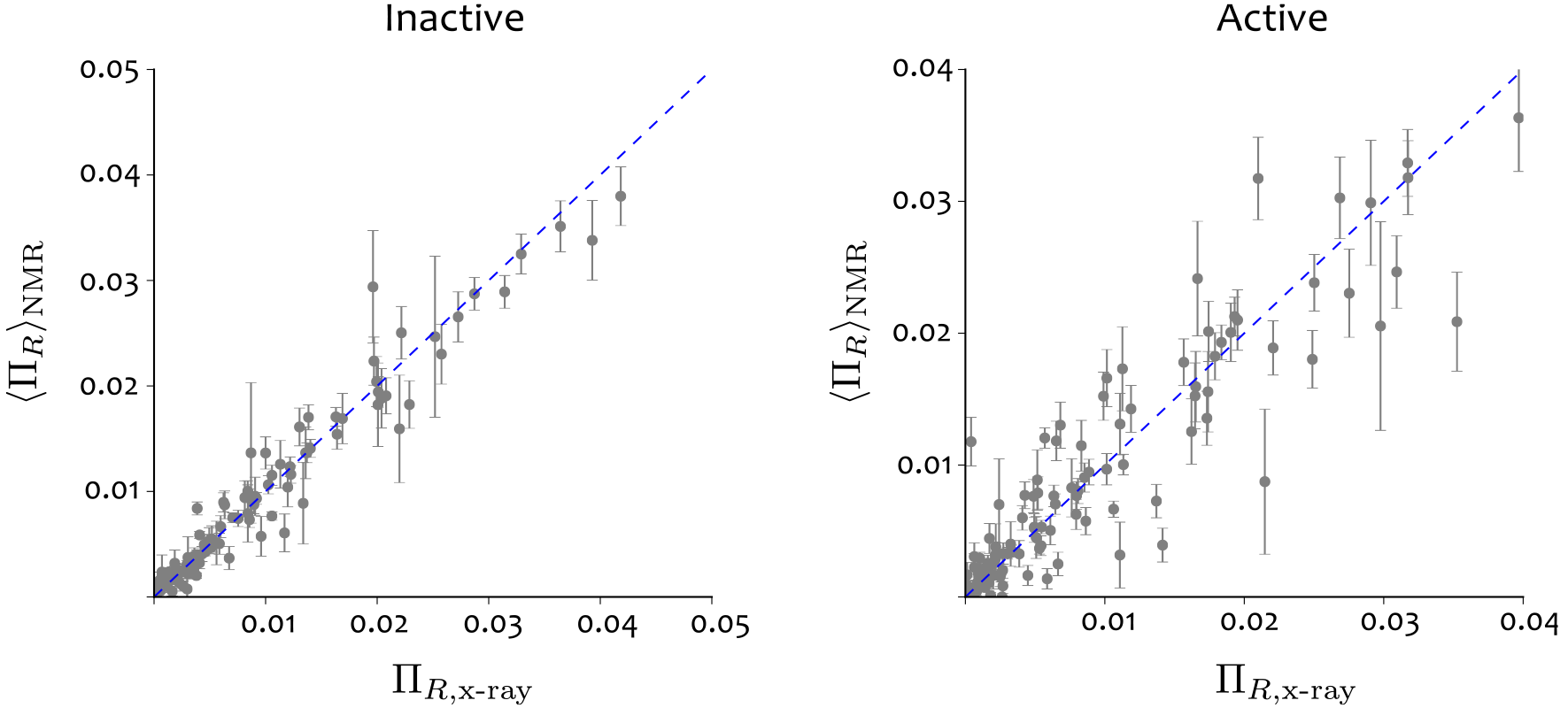
Propensities computed from CheY NMR ensembles. Average propensity obtained from all structures in an NMR ensemble of CheY against the propensity obtained from the corresponding X-ray structure for inactive (left) and active (right). The inactive ensemble contains 20 structures and the active ensemble contains 27 structures. The error bars show the standard deviation of the propensities Π_R_ over the NMR ensemble. Both the variance and the deviation from the X-ray structure is greater for the active conformation.

### Section 4 — The protein reference set from the SCOP database and absolute quantile scores

As discussed in Section IID, we have collected a random reference set of 100 proteins drawn from the Structural Classifiation of Proteins (SCOP) database [13]. This reference protein set is used to obtain
absolute quantile scores for the propensities, as detailed in Materials and Methods (Section IVC). Here we give further details on the reference set and the comparison of absolute and intrinsic quantile scores.

**SCOP database**: Protein domains in the SCOP database are classified according to a hierarchy based on structural similarity. Although proteins are additionally divided into superfamilies and subfamilies according to structural and sequence similarity, the major classes are:

1. All *α* protein domains containing only alpha-helices
2. All β: protein domains containing only beta-sheets
3. Alpha and beta (*α*/*β*): protein domains containing both *α*-helices and *β*-sheets, with mainly parallel *β*-sheets.
4. Alpha and beta (*α* + *β*): protein domains containing both *α*-helices and *β*-sheets, with mainly antiparallel *β*-sheets.
5. Multi-domain: folds of two or more domains from different classes.

We chose 20 proteins from each of these five classes uniformly at random from all proteins in each class, yet choosing only from structures where there is a ligand bound to the active site.

**Absolute quantile scores**: On this set of 100 proteins, we then identified the active site in each protein
and computed the propensity for all its bonds relative to the active site. Across the set of 100 proteins in the reference set, we have a total of 465,409 non-covalent bonds, on which we apply quantile regression to obtain absolute quantile scores *p^ref^*. In Figure S3 below, the quantile scores *p_b_* for all the bonds of the three proteins studied in detail in the main text (caspase-1, Che-Y, h-Ras) are plotted against their absolute quantile score 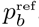, showing a good correlation overall. In general, we observe a tighter correlation for larger proteins (e.g., caspase-1), as a result of the QR fit being based on the number of bonds, *E*, which is related to the size of the protein.

**Figure S3:**
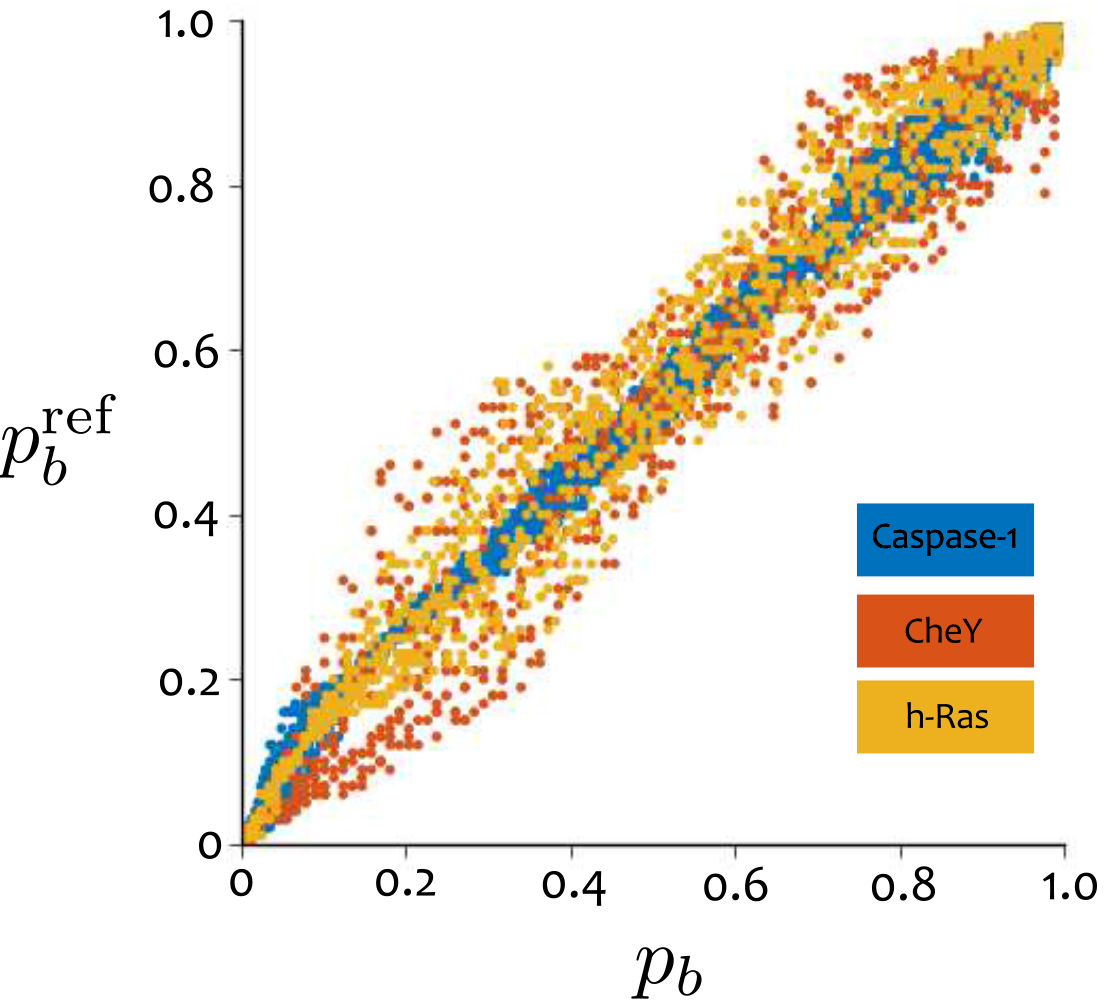
Absolute quantile scores versus intrinsic quantile scores. The absolute quantile scores calculated from the reference set 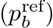 are plotted against the intrinsic quantile scores (*p_b_*) for caspase-1 (blue), CheY (red), and h-Ras (yellow).

### Section 5 — Bond-to-bond propensities of the allosteric test set

**Figure S4:**
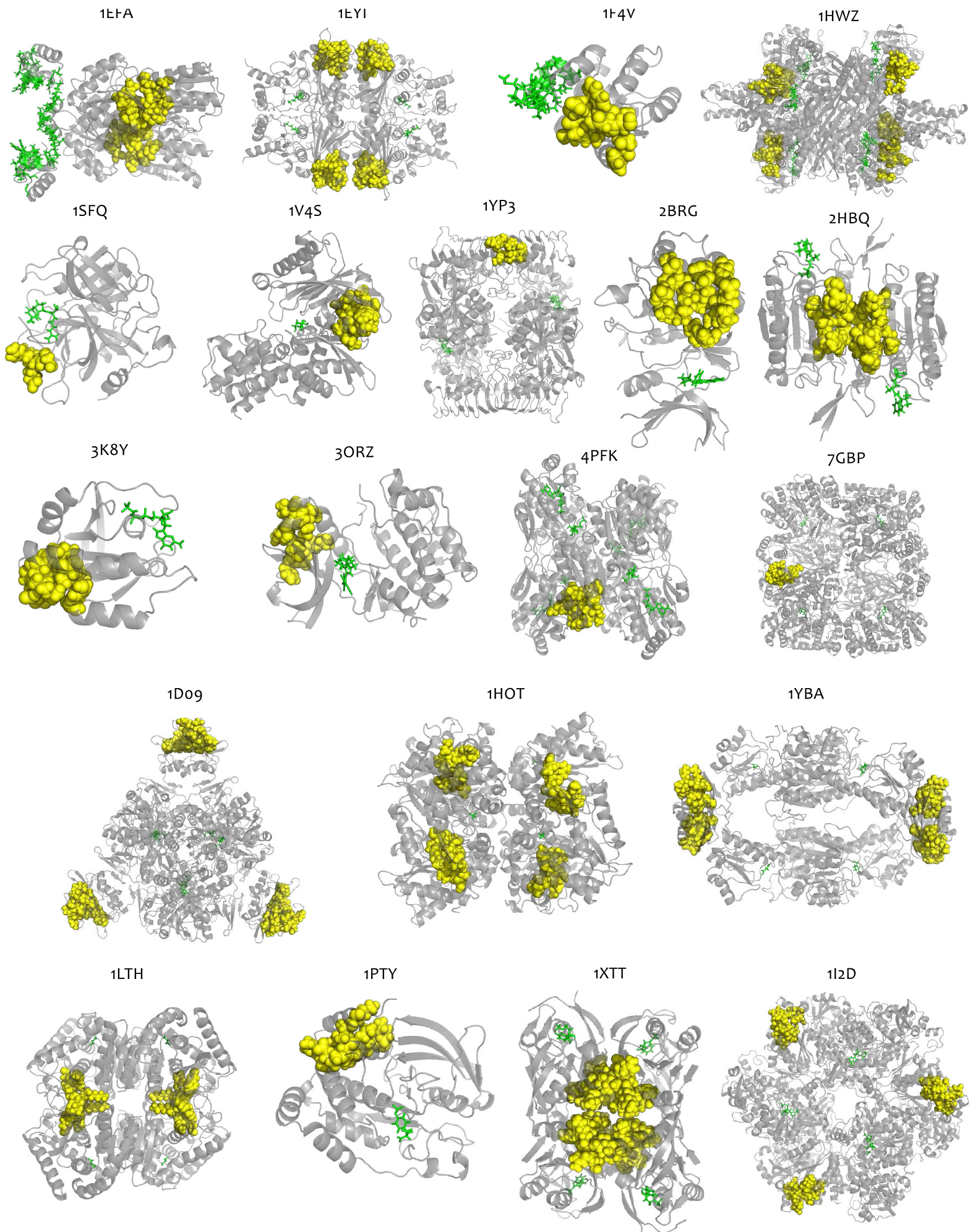
Allosteric test set. The structures of the 20 proteins in the allosteric test set are shown with the active site ligand (green sticks) and allosteric site residues (yellow spheres).

#### 5.1 Description of the allosteric test set

As discussed in the main text (Section IIE), we have constructed a test set of 20 allosteric proteins on which to benchmark our algorithm. Each protein in our test set has a structure with a bound active site ligand and a structure with a bound allosteric ligand. If the protein is allosterically activated then we use a single structure in which the protein is complexed with both the activator and the active site ligand. Ref. [14] collected a test set of 15 allosteric proteins for which both active site bound and allosteric site bound structures are available. We have used 10 of these proteins (the other five were found to be unsuitable for our analysis due to the presence of many non-standard amino-acids, mismatch between the oligomeric state of the active and inactive structures, or the absence of an allosteric ligand). We have enlarged the set with a further 10 proteins from an extensive search of the literature. The structures of the 20 proteins are shown in Figure S4, with the active site indicated by the green ligand, and the allosteric site indicated by the yellow spheres. The allosteric site is defined as any residue containing an atom within 4Åof the allosteric ligand; allosteric site bonds are defined as any weak interactions formed by an allosteric residue. Full details of the proteins and allosteric site residues are shown in Table S2.

**Table S2.**
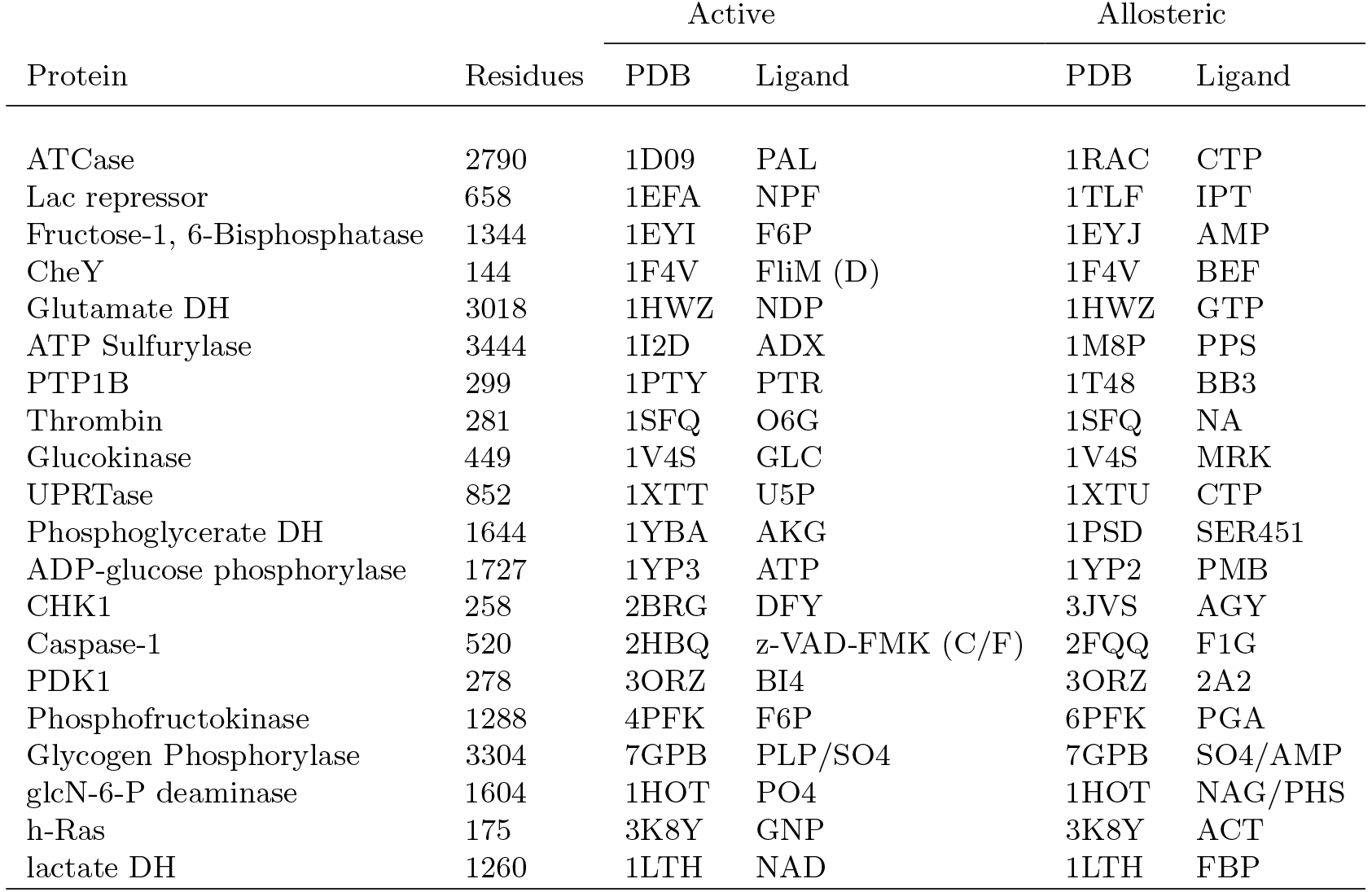
Proteins in the allosteric test set. The active site and allosteric site bound structures for each of the 20 test set proteins. If the protein is allosterically activated then the PDB ID for both states will be the same. The ligand identifier is that used in the PDB file. Exceptions to this are CheY and caspase-1. As the ligand in these proteins is a peptide, the name and chain ID of the peptide is given instead.

#### 5.2 Summary of results on the allosteric test set

As explained in the main text (Section IIE and Materials and Methods, Section IVD), for each of the 20 proteins in the test set, we analyse the propensities of all bonds with respect to the active site of the bound structure, using the ligands shown in Fig. S4 as the source for the bond-to-bond propensity calculations. For each protein, we obtain the propensity Π_*b*_ of every weak bond and its associated quantile score (*p_b_*). To establish their statistical significance, the bond quantile scores *p_b_* (and residue averaged quantile scores *p_R_*) of the allosteric site are compared against an ensemble of randomly generated surrogate sites from each protein. The ensemble of surrogate sites is constructed at random by picking sites that satisfy two structural constraints: (i) they have the same number of residues as the allosteric site; and (ii) their diameter (the maximum distance between any two atoms in the site) is no larger than that of the allosteric site. The sites are generated using Algorithm 1 with pseudocode given below. The propensities averaged over the ensemble of surrogate sites are then used for statistical comparison with the allosteric site. We also obtain absolute propensity scores for each bond 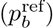 by comparing against the reference SCOP ensemble of 100 proteins. These quantities are defined in the main text (Materials and Methods, Section IVD).

**Algorithm 1.**
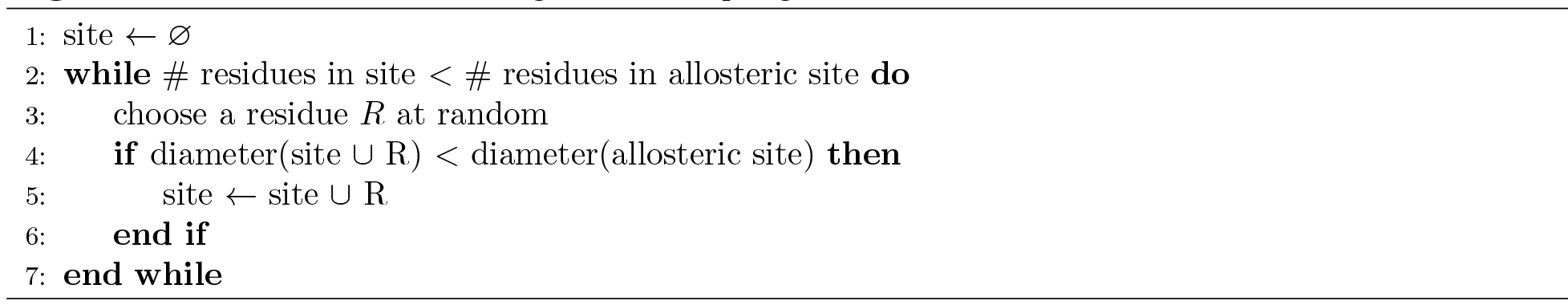
Pseudocode for surrogate site sampling

Using all these scores we obtain our four statistical measures of significance summarised in Table S3. These numerical results are presented also in the form of a graph in Figure 7 of the main text.

**Table S3.**
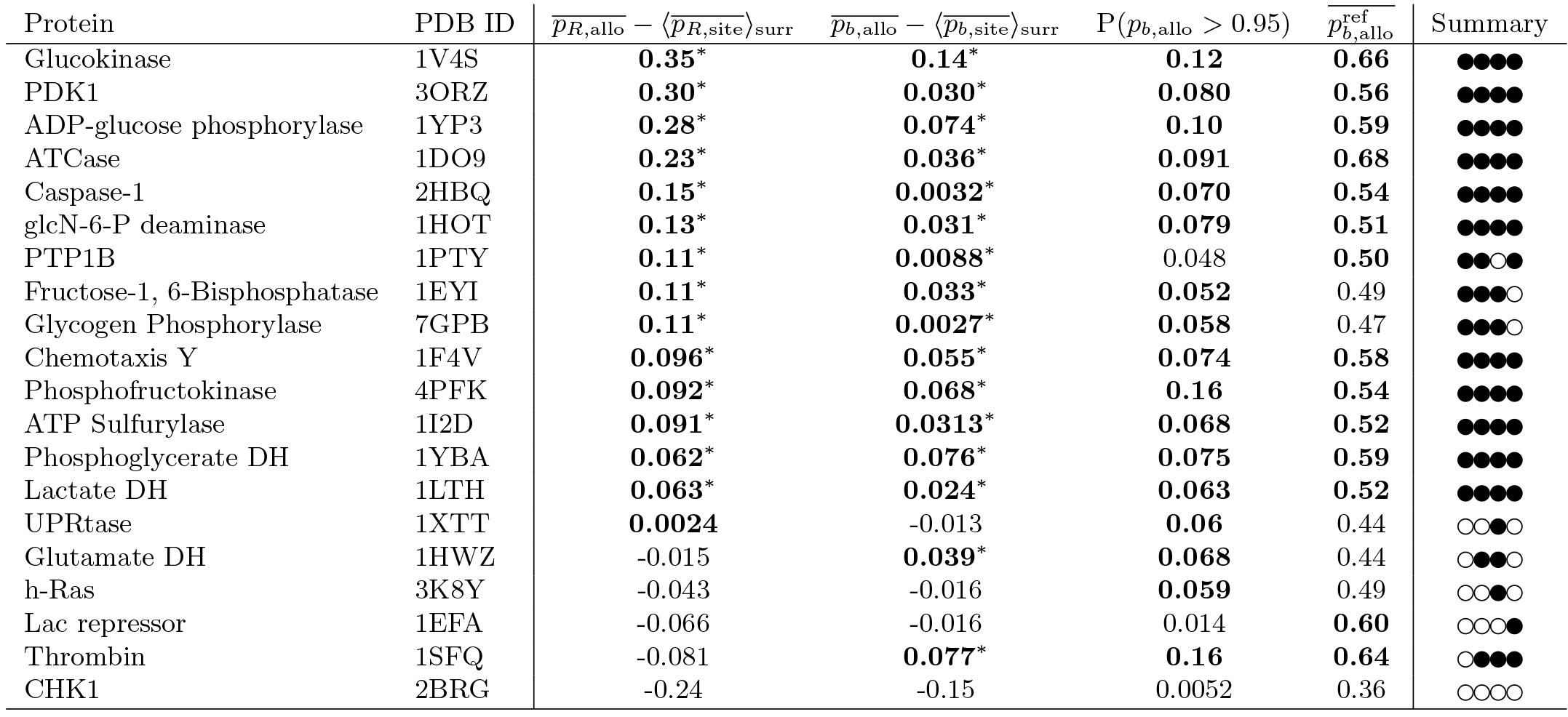
Allosteric site quantile scores in test set proteins. The four scores described in Figure 7 of the main text for the test set of 20 proteins. The difference between the allosteric site average quantile score and the average surrogate site score for both residues and bonds are shown in bold if they are greater than 0, and starred if they lie above the 95% confidence interval computed by a bootstrap with 10000 resamples. The average reference quantile score 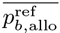 is shown in bold if it is greater than 0.5 (the expected value). The proportion *p_b,allo_* > 0.95 is shown in bold if it is greater than 0.05.

### Section 6 — Robustness of the bond-to-bond propensities to random perturbations of the weak interactions

Proteins are dynamic objects undergoing motions and fluctuations under the influence of the environment. Such dynamic fluctuations induce changes in the bond energies of the protein, potentially leading to the breaking of weak bonds (hydrogen bonds, salt bridges, hydrophobic tethers). As discussed in the main text when studying the NMR ensemble of conformations of CheY (Section IIB.3), whilst there is considerable agreement between the results from the NMR structures and the X-ray structure (Fig. S2), the variability in the ensemble can reveal further information. It is also important to check that the computation of propensities is generally robust to the presence of such noise. To do this, we have developed two schemes to add random perturbations to our protein networks. These schemes mimic the effect of small dynamic fluctuations, without carrying out expensive molecular dynamics simulations.

Firstly, for each of the 20 proteins in our dataset, we add zero mean Gaussian noise to the edge weights (energies) of non-covalent bonds in the graph, so as to mimic the effect of thermal fluctuations. Note that we allow the bonds to break if their randomised energy becomes zero. We then recompute our quantile scores for the allosteric site for 10 realisations of the noisy networks generated after the addition of the Gaussian fluctuations. We do this for 3 levels of noise, i.e., we increase the standard deviation of the Gaussian from 1kT=0.6 kcal/mol to 4kT=2.4 kcal/mol. The average results of these randomisations for all proteins in the allosteric test set are presented in Table S4. Our calculations show that the results are generally robust to fluctuations induced in this way: the signal at the allosteric site only drops slightly when introducing relatively high levels of noise.

**Table S4.** Robustness of propensity scores to additive randomness. Mean (± standard deviation) of propensity scores 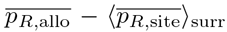 computed from randomisations of the protein networks of the allosteric test set obtained by adding Gaussian noise to the edge weights (bond energies). The noise level varies between 1kT and 4kT (corresponding to the standard deviation of the added Gaussian) and at each noise level the results were calculated from 10 randomised graphs. The difference between the allosteric site average quantile score and the average surrogate site score for both residues and bonds are shown in bold if they are greater than 0, and starred if they lie above the 95% confidence interval computed by a bootstrap with 10000 resamples. The unperturbed result is also shown for comparison.

| PDB ID | Unperturbed network | Gaussian noise 1kT | Gaussian noise 2kT | Gaussian noise 4kT |
| --- | --- | --- | --- | --- |
| 1V4S | **0.35\*** | **0.32±0.011\*** | **0.31±0.019\*** | **0.27±0.017\*** |
| 3ORZ | **0.30\*** | **0.28±0.0087\*** | **0.23±0.0090\*** | **0.24±0.014\*** |
| 1YP3 | **0.28\*** | **0.27±0.0010\*** | **0.25±0.0088\*** | **0.17±0.016\*** |
| 1D09 | **0.23\*** | **0.22±0.0071\*** | **0.21±0.0024\*** | **0.20±0.0035\*** |
| 2HBQ | **0.15\*** | **0.18±0.0096\*** | **0.20±0.0058\*** | **0.13±0.0097\*** |
| 1HOT | **0.13\*** | **0.13±0.0061\*** | **0.098±0.024\*** | **0.12±0.021\*** |
| 1PTY | **0.11\*** | **0.096±0.020\*** | **0.11±0.022\*** | **0.088±0.031\*** |
| 1EYI | **0.11\*** | **0.13±0.0065\*** | **0.13±0.0022\*** | **0.16±0.0050\*** |
| 7GPB | **0.11\*** | **0.096±0.018\*** | **0.13±0.010\*** | **0.14±0.015\*** |
| 1F4V | **0.096\*** | **0.093±0.018\*** | **0.14±0.0097\*** | **0.12±0.027\*** |
| 4PFK | **0.092\*** | **0.091±0.0052\*** | **0.11±0.022\*** | **0.12±0.0075\*** |
| 1I2D | **0.091\*** | **0.14±0.029\*** | **0.14±0.030\*** | **0.14±0.037\*** |
| 1YBA | **0.062\*** | **0.076±0.0034\*** | **0.091±0.0048\*** | **0.073±0.0051\*** |
| 1LTH | **0.063\*** | **0.070±0.0099\*** | **0.063±0.016\*** | **0.039±0.019\*** |
| 1XTT | **0.0024** | **0.0084±0.0069** | **0.015±0.0083** | **0.0077±0.0070** |
| 1HWZ | -0.015 | -0.0090±0.0071 | -0.0028±0.0043 | **0.011±0.0065** |
| 3K8Y | -0.043 | -0.033±0.012 | -0.012±0.010 | -0.025±0.022 |
| 1EFA | -0.066 | -0.047±0.0054 | -0.019±0.0078 | -0.0027±0.0077 |
| 1SFQ | -0.081 | -0.090±0.0089 | -0.083±0.023 | -0.10±0.028 |
| 2BRG | -0.24 | -0.23±0.010 | -0.24±0.013 | -0.23±0.026 |

Secondly, to test a different kind of variability introduced by the environment, we have considered the effect of breaking all bonds in our network with energy below a threshold. Starting with the original unperturbed structure, all weak bonds below a given threshold are removed from the graph. In this way, we mimic the possibility of extended structural changes that could lead to breaking of bonds in a more global fashion.

For each of the 20 proteins in the test set, we generate two perturbed networks obtained by bond removal of *all* bonds with energy below two different thresholds: 0.5 kT≃ 0.3 kcal/mol and 1kT ≃ 0.6 kcal/mol. The effect of this thresholding is extensive. For the 0.5kT threshold, we delete all hydrophobic tethers and electrostatic interactions as well as a percentage of hydrogen bonds that ranges from 31% in 1SFQ to 44% in 1HWZ and 1LTH. For the 1kT threshold, even further hydrogen bonds are removed, corresponding to eliminating 44% of H-bonds in 1SFQ up to 57% of the H-bonds in 7GPB, 2BRG, 1LTH (in addition to all hydrophobic interactions).

The calculations of the propensity for the thresholded networks for all 20 proteins in our test set are presented in Table S5. Our results show that, overall, the propensity of the allosteric site remains largely robust to such changes across all 20 proteins considered, yet with notable differences in the magnitude of the effect across the set. In some proteins, the signal at the allosteric site is mildly affected by bond deletion (e.g. 3ORZ, 1YP3, 2HBQ, 1HOT, 1PTY). In other cases, however, the deletion of weaker hydrogen bonds has a large effect in destroying the communication between the allosteric site and the active site (e.g. 1V4S, 1D09, 1EYI, 7GPB, 1F4V). These differences could be a measure of how robust the allosteric signalling is to energetic fluctuations in the local environment of the protein, and also provide clues as to different structural features connected with the distributed nature of allosteric signalling in the different proteins. The study of such differences will be the object of future work.

**Table S5.** Robustness of propensity scores to deletion of weak bonds. The propensity score 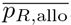 — 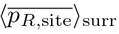 for networks obtained by deleting all bonds below two energy thresholds. The results are shown in bold when they are greater than 0 and starred if they lie above the 95% confidence interval computed by a bootstrap with 10000 resamples. The unperturbed score is reported also for comparison.

| PDB ID | Unperturbed network | Threshold 0.5 kT | Threshold 1kT |
| --- | --- | --- | --- |
| 1V4S | **0.35\*** | **0.061\*** | **0.049\*** |
| 3ORZ | **0.30\*** | **0.24\*** | **0.25\*** |
| 1YP3 | **0.28\*** | **0.24\*** | **0.30\*** |
| 1D09 | **0.23\*** | **0.088\*** | **0.10\*** |
| 2HBQ | **0.15\*** | **0.16\*** | **0.18\*** |
| 1HOT | **0.13\*** | **0.14\*** | **0.17\*** |
| 1PTY | **0.11\*** | **0.13\*** | **0.080\*** |
| 1EYI | **0.11\*** | **0.026\*** | **0.022\*** |
| 7GPB | **0.11\*** | **0.056\*** | **0.062\*** |
| 1F4V | **0.096\*** | -0.0010 | **0.0085\*** |
| 4PFK | **0.092\*** | **0.17\*** | **0.20\*** |
| 1I2D | **0.091\*** | **0.0012** | -0.033 |
| 1YBA | **0.062\*** | **0.079\*** | **0.052\*** |
| 1LTH | **0.063\*** | **0.056\*** | -0.081 |
| 1XTT | **0.0024** | -0.016 | -0.023 |
| 1HWZ | -0.075 | -0.016 | -0.20 |
| 3K8Y | -0.043 | -0.14 | -0.16 |
| 1EFA | -0.066 | **0.052\*** | **0.051\*** |
| 1SFQ | -0.081 | **0.073\*** | **0.11\*** |
| 2BRG | -0.24 | -0.20 | -0.17 |

### Section 7 — Propensities from residue-residue interaction networks

The computational efficiency of our methodology allows us to analyse all-atom networks without many of the restrictions on system size inherent to other methods. Proteins or protein complexes of hundreds of thousands of atoms can be analysed in a few minutes on a standard desktop. We can thus keep atomistic detail at the single bond level without restricting the scope of the analysis. Hence there is a less acute need to seek computational savings by obtaining coarse-grained representations of proteins at the level of residue interactions. However, it is still instructive to consider propensity measures computed from residue-level networks (RRINs) [15]. We have undertaken this comparison for all 20 proteins in our test set and report the results below.

As discussed in the main text (Sections IIA and IIB), in some cases (e.g., caspase-1, Fig 1b) we found that the additional information contained in the atomistic network leads to increased signal in the detection of the allosteric site, whereas in other cases (e.g., CheY), RRINs already capture well the site connectivity that reveals the presence of the allosteric site. Our analysis of the full test set (Table S6) confirms that the results from RRINs depend on the protein analysed, and also vary substantially depending on the choice of the cut-off distance (a tunable parameter which needs to be chosen when generating the coarse-grained RRINs).

The coarse-grained RRINs for each of the 20 proteins in the test set were obtained by submitting the corresponding PDB files to the oGNM server [16]. We obtained RRINs at four different cut-off radii: 6 Å, 7 Å, 8 Å and 10 Å. The cut-off radius is a tunable parameter necessary to generate a RRIN from PDB files, which establishes how close two residues must be in order to be connected in the RRIN. A range of different cut-off radii has been used throughout the literature. However, the usual radius is around 6.7-7.0 Å, which corresponds to the first coordination shell [17].

Table S6 shows the propensity score of the allosteric site 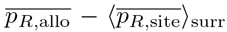, computed from RRINs obtained at four cut-offs (between 6Å and 10Å) for the 20 proteins in the allosteric test set. For comparison purposes, we also report the same score obtained from the all-atom network. It is important to note that this is just one of four scores obtained from the all-atom network, reflecting only the *averaged* behaviour over the residues. This score is complemented by the three other bond-based statistics, which can pick up inhomogeneities in the propensities of the bonds in the allosteric site, as given by the All-atom Summary column carried over from Table S3.

Our results indicate broad consistency between RRINs and the all-atom network. However, the RRIN results vary widely depending on the choice of cut-off radius in the generation of the network. Moreover this variability with respect to the cut-off behaves differently for each of the proteins. As an illustration, the allosteric site of caspase-1 (2HBQ) was not found to be significant in the RRINs with cut-off radii of 6 Å, 7 Å and 8 Å and only weakly significant for 10 Å, whereas 1LTH and 2BRG are both only detected in RRINS with cut-off radius of 6 Å but not for larger radii. Our results are consistent with previous studies that found that allosteric pathway identification in RRINs is dependent on the chosen cut-off [18]. For the different cut-offs, the number of proteins with *p*_*R*,allo_ > *p*_*R*,rest_ varies between 11/20 (at 7, 8, and 10Å) and 13/20 (at 6Å), and only 8/20 proteins have *p*_*R*,allo_ > *p*_*R*,rest_ for the RRINS at *all* the cut-off radii. This is compared to 15/20 proteins for the atomistic network.

Even when the allosteric site is detected in the RRIN, the signal when using the atomistic network is considerably higher in a number of proteins (e.g., 1V4S, 1YP3, 7GPB, 1I2D, 2HBQ). In other cases (e.g., 1EYI, 4PFK), the RRIN directly loses the detectability of the allosteric site even if the cut-off is adjusted. This observation suggests that these are proteins where the specific chemistry of intra-protein bonds is important for the allosteric communication.

On the other hand, there are several other cases (e.g., 3ORZ, 1D09, 1HOT, 1PTY, 1LTH) where the RRIN can provide similar results to the atomistic network, yet still with some variability depending on the choice of appropriate cut-off. Interestingly, there are also some proteins (specifically 1F4V, 1YBA, 3K8Y and 2BRG) in which the propensity score is higher for RRINs than for the atomistic network. In these cases, there tends to be a large heterogeneity in the propensities of the bonds in the allosteric site (see Figure 7 in the main text) with some bonds with large negative values as well as other bonds with large positive values. Our bond statistical measures can account for some of this variability. Indeed, both 1F4V and 1YBA are detected by all our four bond measures, and 3K8Y is picked by the measure based on the distributions of *p_b_*. Intriguingly, only 2BRG (corresponding to CHK1) cannot be detected by our bond measures. This suggests other areas of future research, in which the importance of averaging at the level of pathways could be used to enrich the findings presented here.

**Figure S5:**
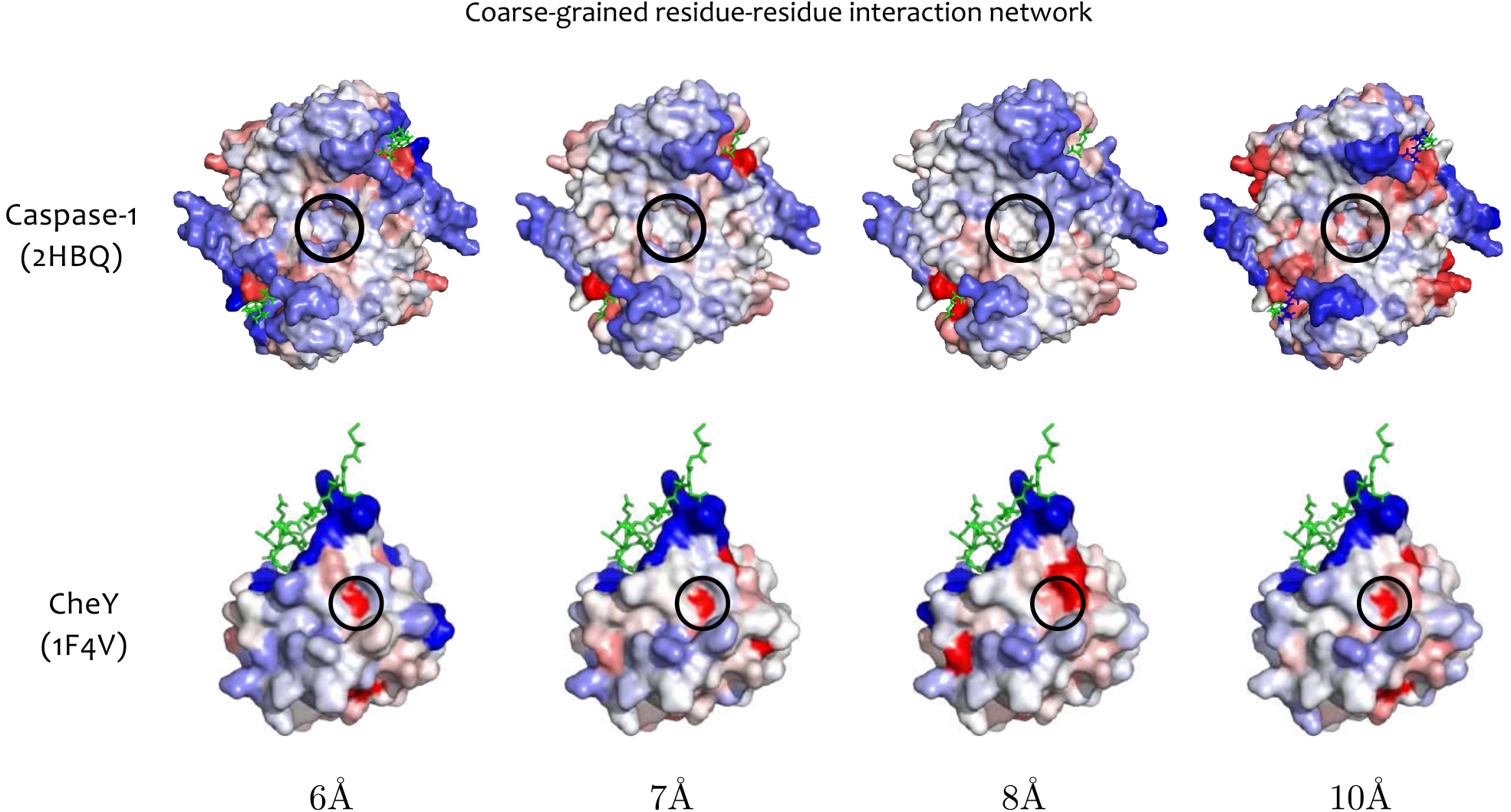
Quantile scores computed from RRINs for caspase-1 and CheY at different cut-off radii. Surface mapping of the residue quantile scores *p_R_* of caspase-1 and CheY for RRINs generated with radii cut-offs from 6 Å and 10 Å. The active-site ligand is shown in green sticks and the allosteric site is circled. The allosteric site in caspase-1 is not identified for 6, 7, and 8 Å. It is identified at 10 A, but the signal is weaker than when using an atomistic graph. In contrast, for CheY the allosteric site is identified as significant across the full range of cut-offs.

**Table S6.**
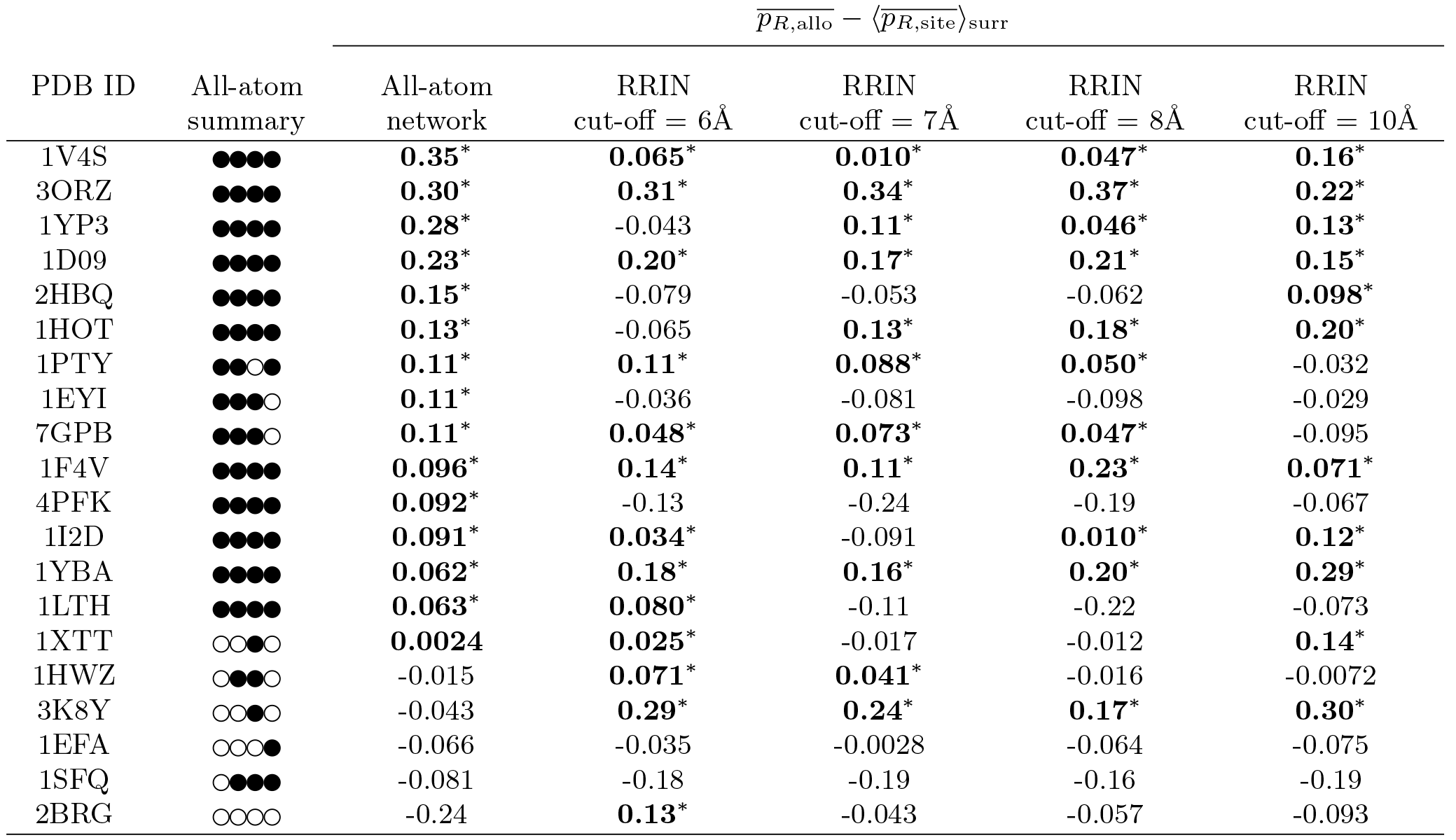
Propensities computed from RRINs. Values of 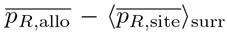 for residue-residue interaction networks with four cut-off radii from 6Å-10Å. The propensity scores are shown in bold if they are greater than 0, and starred if they lie above the 95% confidence interval computed by a bootstrap with 10000 resamples. The comparable statistic computed from the all-atom network is also presented, as well as the summary of the four bond statistics for each protein from Table S3.

